# Neuron-glia interaction through Serotonin-BDNF-NGFR axis enables regenerative neurogenesis in Alzheimer’s model of adult zebrafish brain

**DOI:** 10.1101/748970

**Authors:** Prabesh Bhattarai, Mehmet Ilyas Cosacak, Violeta Mashkaryan, Sevgican Yilmaz, Stanislava Dimitrova Popova, Nambirajan Govindarajan, Kerstin Brandt, Yixin Zhang, Caghan Kizil

## Abstract

It was recently suggested that supplying the brain with new neurons could counteract Alzheimer’s disease. This provocative idea requires further testing in experimental models where the molecular basis of disease-induced neuronal regeneration could be investigated. We previously found that zebrafish stimulates neural stem cell (NSC) plasticity and neurogenesis in Alzheimer’s disease and could help to understand the mechanisms to be harnessed for develop new neurons in diseased mammalian brains. Here, by performing single-cell transcriptomics, we found that Amyloid toxicity-induced Interleukin-4 induces NSC proliferation and neurogenesis by suppressing the tryptophan metabolism and reducing the production of Serotonin. NSC proliferation was suppressed by Serotonin via downregulation of BDNF-expression in Serotonin-responsive periventricular neurons. BDNF enhances NSC plasticity and neurogenesis via NGFRA/NFkB signaling in zebrafish but not in rodents. Collectively, our results suggest a complex neuron-glia interaction that regulates regenerative neurogenesis after Alzheimer’s disease conditions in zebrafish.

**Key findings:**

- Amyloid-induced Interleukin-4 suppresses Serotonin (5-HT) production in adult zebrafish brain
- 5-HT affects *htr1*-expresing neurons and suppresses *bdnf* expression
- BDNF enhances plasticity in neural stem cells via NGFRA/NFkB signaling
- BDNF/NGFRA signaling is a neuro-regenerative mechanism in zebrafish but not in mammals.

## Introduction

Alzheimer’s disease (AD) entails versatile pathological changes such as synaptic degeneration, neuronal death, chronic inflammation, impaired vasculature function and reduced plasticity of neural stem cells (De Strooper and Karran, 2016; Moreno-Jimenez et al., 2019; Selkoe and Hardy, 2016). The cognitive decline that is observed in AD patients and experimental animal models is mainly caused by the reduced neural network integrity (Arendt, 2009). The efforts to rescue the cognitive decline and neuronal death traditionally largely focused on the neuronal compartment and aimed at preventing the death of the neurons while this has not yielded in desired success in clinics (Cummings et al., 2016; Mehta et al., 2017). An alternative approach was suggested to be to complement the treatment in neuronal compartments with increasing the production of new neurons to provide resilience and strength to the diseased circuitry (Choi et al., 2018; Moreno-Jimenez et al., 2019; Mu and Gage, 2011; Tincer et al., 2016). Yet, neurogenesis in human brains is quite controversial (Arellano et al., 2018; Cipriani et al., 2018; Dennis et al., 2016; Duque and Spector, 2019; Sorrells et al., 2018). Although many reports documented the presence of adult neurogenesis in human brains (Boldrini et al., 2018; Ernst et al., 2014; Kempermann et al., 2018; Magnusson and Frisen, 2016; Moreno-Jimenez et al., 2019; Spalding et al., 2013), and several studies demonstrated that boosting the neurogenesis might be a viable option for alleviating the cognitive decline (Casse et al., 2018; Choi et al., 2018; Martinez-Canabal, 2014; Papadimitriou et al., 2018; Rodriguez and Verkhratsky, 2011), the potential benefits of neurogenic outcome in AD conditions requires further investigation and critical testing. Additionally, in AD conditions, mammalian NSCs reduce the proliferative ability dramatically, and for neurogenesis to become a viable option for treatment of neurological disorders, mammalian NSCs must become plastic first. Therefore, examining how NSCs could be made proliferative and neurogenic during the course of AD could provide important clinical ramifications towards the treatment of this disease (Kizil and Bhattarai, 2018a); however, little is known about the mechanisms by which neural stem cells would enhance their proliferative response (Dong et al., 2019; Kempermann et al., 2018; Morrens et al., 2012). Recently, we established a zebrafish model, which can recapitulate the symptoms of AD in humans such as neuronal death, synaptic degeneration, chronic inflammation and cognitive decline (Bhattarai et al., 2017a; Bhattarai et al., 2016). Interestingly, in contrast to humans, zebrafish brain could enhance NSC proliferation and neurogenesis through a previously unidentified neuro-immune regulation involving Interleukin-4 (IL4). IL4 secreted by dying neurons and activated microglia, which in turn activated NSC proliferation (Bhattarai et al., 2016). We also found that IL4 could revert the pathological effects on NSC in an in vitro 3D reductionist model of AD (Papadimitriou et al., 2018). Yet, the mechanisms by which IL4 regulates NSC proliferation and neurogenesis after Amyloid toxicity remained unknown. In this manuscript, using single cell sequencing, we identified a previously undocumented IL4-dependent mechanism that regulates NSC plasticity in adult zebrafish brain, where IL4 regulates production of Serotonin, which suppresses production of the brain-derived neurotrophic factor (BDNF) in periventricular neurons juxtaposing the NSCs. We found that BDNF is required to activate NSC plasticity, proliferation of NSCs, and neurogenesis through its receptor nerve growth factor receptor A (NGFRA), the blockage of which reduces NSC proliferation. Overall, our results identify a mechanism by which IL4 regulates NSC plasticity through Serotonin-dependent expression of BDNF in neurons in zebrafish brain, and demonstrate functional heterogeneity of NSCs based on receptor expression. Our results will provide a conceptual basis for neuro-immune regulation of NSCs, neuronal control of NSC proliferation, and differential response of NSC subtypes to various signals in zebrafish. Such understanding could be instrumental in the efforts to develop novel therapies for Alzheimer’s disease through increased neurogenesis.

## Results

To determine the effects of Interleukin-4 (IL4) on neural stem cell (NSC) plasticity in adult zebrafish brain, we administered IL4 through cerebroventricular microinjection, dissected the telencephalon and performed a whole transcriptome profiling in control and IL4-injected brains (Figure 1A, Figure S1). By comparing the gene expression profiles, we found that IL4 administration increased the expression of 285 genes while downregulated 1435 genes (Figure 1B, Dataset S1). To determine the pathways affected by IL4, we performed KEGG analyses and observed that one of the significantly regulated pathways was Tryptophan metabolism (Figure 1C, Figure S1). Specifically, we observed that the enzymes generating Serotonin from Tryptophan were downregulated (Figure 1D, Dataset S2), suggesting that IL4 might reduce the production of Serotonin (5-HT). To test whether Amyloid-beta42 (Aβ42, which induces IL4 expression (Bhattarai et al., 2016)) and ectopic IL4 would reduce the availability of 5-HT in adult zebrafish brain, we performed immunohistochemistry for 5-HT (Figure 1E) and observed that Aβ42 and IL4 significantly reduces the 5-HT immunoreactivity in adult zebrafish telencephalon (Figure 1F) but the cell bodies at more caudal regions still existed (Figure S2A-C), suggesting that Aβ42/IL4 suppresses Serotonin production rather than the innervation. Additionally, the effects of IL4 is specific to Serotonergic system, as in our deep sequencing analyses we did not observe changes in KEGG analyses in other neurotransmitter pathways such as dopamine, histamine or noradrenalin (Dataset S2).

**Figure1:**
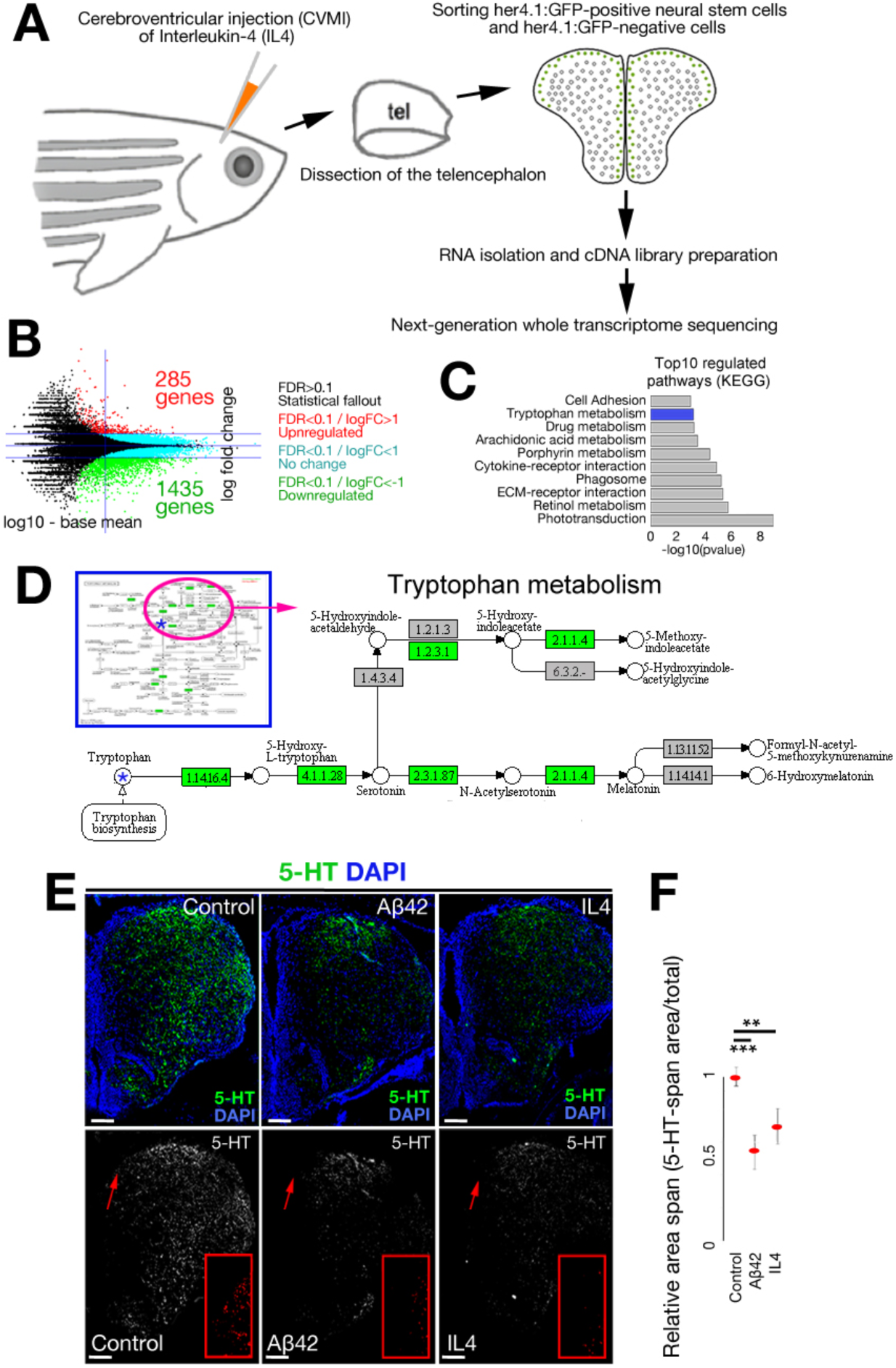
Interleukin-4 regulates Tryptophan metabolism. (A) Schematic view of the experimental pipeline for whole transcriptome sequencing after Interleukin-4 (IL4) treatment. (B) MA-plot for differentially expressed genes (DEGs). (C) GO-term analyses on DEGs. (D) Modified KEGG-pathway view of Tryptophan metabolism. Green indicates the enzymes downregulated by IL4. (E) Immunohistochemistry (IHC) for Serotonin (5-HT) in control (left), Aβ42-injected (middle), and IL4-injected (right) brains. Single channel images show 5-HT. Red insets are high-magnification images of arrowed regions. (F) Quantification of 5-HT-span area density under the conditions of F. n = 3 animals for experiments. Scale bars equal 100 µM. Data are represented as mean ± SEM. See also Figure S1.

Since Aβ42/IL4 enhances neural stem cell (NSC) proliferation in adult zebrafish brain (Bhattarai et al., 2017a; Bhattarai et al., 2016; Bhattarai et al., 2017b; Kizil, 2018; Kizil and Bhattarai, 2018b) and they downregulate 5-HT (Figure 1E,F), we hypothesized that 5-HT could be negatively affecting NSC plasticity. To test this, we injected 5-HT into adult zebrafish brain and analyzed the proliferation of NSCs (S100b/PCNA double positive cells) (Figure 2A-C). We observed a significant reduction of NSC proliferation after 5-HT injection (Figure 2C). This reduction has implications in neurogenesis from NSCs, as the analysis of BrdU-labeled newborn neurons at 14 days after injection of PBS (Figure 2D) or 5-HT (Figure 2E) showed that 5-HT reduces the neurogenic outcome (Figure 2F). To verify our results, we used a transgenic reporter line that marks NSCs with GFP driven under the *her4.1* promoter and injected Aβ42, IL4 or 5-HT followed by immunohistochemisty for GFP and PCNA (Figure 2G). Consistent with our previous results, Aβ42/IL4 injection increased NSC proliferation, while 5-HT reduced the number of proliferating progenitors (Figure 2H), indicating that 5-HT has a negative impact on NSC plasticity in zebrafish telencephalon. To further test whether the negative effects of 5-HT would be reversed by Aβ42 or IL4, we co-injected 5-HT with Aβ42 and IL4, and observed that the reduction in NSC proliferation by 5-HT can be abrogated by Aβ42 or IL4 (Figure S2D-H) indicating that 5-HT signaling acts antagonistically to IL4 in NSCs.

**Figure2:**
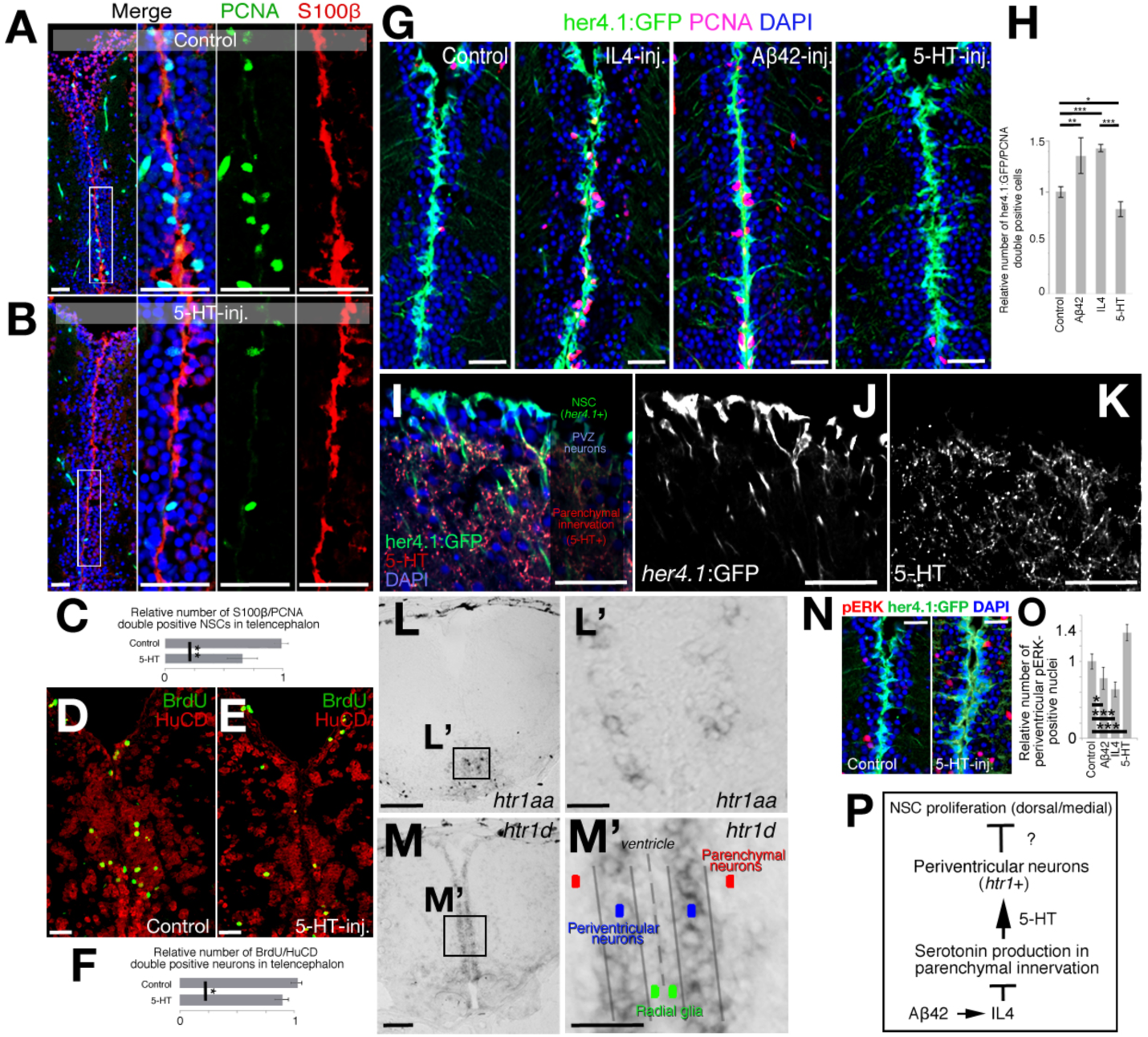
Serotonin regulates neural stem cell plasticity indirectly through HTR1 signaling in periventiruclar neurons. (A, B) Immunohistochemisty (IHC) for S100β and PCNA in control (A) and 5-HT-injected (B) brains. (C) Quantification of proliferating glia in conditions of A and B. (D,E) IHC for BrdU and HuC/D for newborn neurons at 14dpi after BrdU treatment at 2 and 3 days after control (D) and BDNF-injection (E). (F) Quantification of newborn neurons. (G) IHC for GFP (driven by glial promoter her4.1) and PCNA in control, IL4-injected, Aβ42-injected and 5-HT-injected brains. (H) Quantification of proliferating glia in conditions of G. (I-K) IHC for her4.1-driven GFP and 5-HT. The composite image (I) and single fluorescent channels for her4.1:GFP (J) and 5-HT (K). (L) In situ hybridization (ISH) for *htr1a* (L’: close-up image of framed region in L). (M) ISH for *htr1d* (M’: close-up image of framed region in M). (N) IHC for pERK and her4.1-driven GFP in control and 5-HT-injected brains. (O) Quantification of pERK-positive periventricular neurons. (P) Working hypothesis on the indirect regulation of 5-HT on neural stem cell plasticity. n = 4 animals for experiments. Scale bars equal 100 µM. Data are represented as mean ± SEM. See also Figure S2.

Our previous results indicated that a subset of NSCs express IL4 receptor and can therefore be directly regulated by IL4 (Bhattarai et al., 2016; Cosacak et al., 2019). Therefore, we aimed to investigate if 5-HT would affect the same subset of NSCs or whether it affects a distinct population. To determine whether or not 5-HT has a direct effect on NSCs, we determined the spatial organization of NSCs (*her4.1*:GFP, Figure 2I,J) and 5-HT innervation (Figure 2I,K). We observed that NSCs are located apically and are separated by periventricular zone (PVZ) neurons before the proximal front of the Serotonergic innervation in the parenchyma (Figure 2I), suggesting that the effect of 5-HT on NSCs could be indirect. If this hypothesis would be true, NSCs would not express 5-HT receptors. To determine which cells expressed 5-HT receptors, we performed in situ hybridization for 5-HT receptors (*htr* genes) and observed that among 7 serotonin receptor genes and in total 21 isoforms of those genes (Dataset S2), only *htr1a* and *htr1d* gave in situ hybridization signals in adult zebrafish brain (Figure 2L-M’), and serotonin receptor genes are not expressed in the progenitor cells (Figure S2I), supporting previous findings (Norton et al., 2008). *htr1a* was expressed only in ventral regions (Figure 2L,L’) while *htr1d* was present in periventricular region immediately adjacent to the ventricular zone containing the NSCs that span the medial and dorsal regions of the telencephalon (Figure 2M,M’). These findings suggested that the effect of 5-HT on NSCs is not direct, and it could be mediated through periventricular neurons expressing *htr1d* (Figure 2N). If 5-HT would act on periventricular neurons, 5-HT would activate its downstream effector pERK (Huang and Reichardt, 2003) in periventricular regions. Indeed, we found that compared to controls where nuclear pERK is in few cells at the PVZ (Figure 2O), 5-HT injection increases the number of pERK+ nuclei only in the PVZ region (Figure 2N). Quantification of pERK-positive nuclei also confirmed that 5-HT increases pERK-positive nuclei on the contrary to Aβ42/IL4 (Figure 2O), indicating that 5-HT directly affects periventricular neurons but not the NSCs (Figure 2N).

To determine the mechanistic link between 5-HT and NSC plasticity through *hrt1*+ cells (Figure 2P), we performed single cell transciptomics in control and 5-HT-treated adult zebrafish telencephalon by unbiased clustering, determination of cell types and differential gene expression analyses using the methodology we have recently developed (Cosacak et al., 2019; Cosacak et al., 2018) (FigureS3A; Dataset S3). After quality control analyses (Figure S3B), we obtained tSNE clusters, which dissolve into four major cell types: neurons (*eno2, gap43, map1aa, sypb*-positive), glia (*gfap, her4.1, msi1, s100b*-positive); oligodendrocytes (*aplnrb, olig2*-positive) and immune cells (*pfn, lcp1*-positive) (Figure S3C-E). Heat maps reveal marker genes expressed predominantly in those cells (Figure S3C) To determine the cells that are responsive to 5-HT, we plotted the cells expressing *htr1* and observed that *htr1* expression was confined to neurons (Figure S3F). This expression pattern was consistent with in situ hybridization results of *htr1d* (Figure 2M) and was distinct than *il4r.1* expressing cells that were exclusively glia and immune cells ((Bhattarai et al., 2016) and Figure S3F).

To determine the mechanism by which 5-HT-responsive cells affect glia proliferation, we devised an analysis pipeline where we dissected the *htr1*+ cells from the rest and performed differential expression analysis between control and 5-HT-treated brains (Figure 3A; Figure S3G). We found that 349 genes changed their expression levels in *htr1*+ cells after 5-HT treatment (Figure 3B, Dataset S4). We hypothesized that a possible regulation between *htr1*+ neurons and NSCs could be through a paracrine ligand-receptor crosstalk. We found that in *htr1*+ cells, 29 ligands change their expression levels (Figure 3C), five of which also change their expression reciprocally after 5-HT or IL4 treatment (Figure 3D, and data from (Cosacak et al., 2019)). The highest fold change was observed for *bdnf*, which is enhanced by IL4 and downregulated by 5-HT (Figure 3D).

**Figure3:**
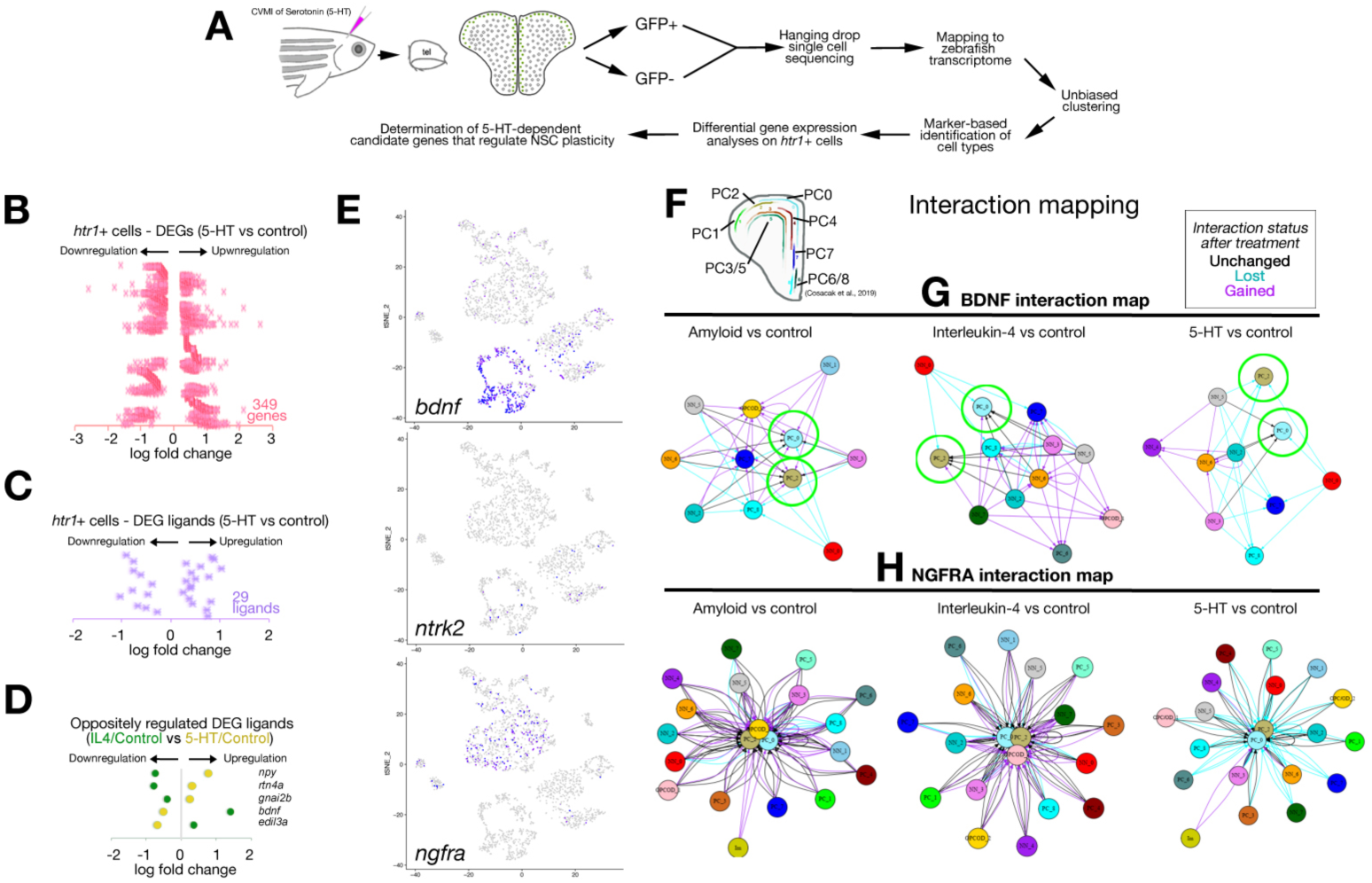
Single cell sequencing after Serotonin treatment in adult zebrafish brain. (A) Schematic workflow for single cell sequencing. (B) Distribution plot for differentially expressed genes (DEGs) in *htr1*-expressing cells after 5-HT treatment. (C) Ligands selected from B. (D) Plots for ligands that change oppositely in 5-HT and IL4 treatment. (E) Feature plots for *bdnf* and its receptors *ntrk2* and *ngfra*. (F) Spatial map of neural stem/progenitor cells (PCs) in adult zebrafish brain as previously described in Cosacak et al., 2019. (G) In silico interaction map for BDNF in Amyloid vs control, IL4 vs control and 5-HT vs control comparisons. (H) In silico interaction map for NGFRA in Amyloid vs control, IL4 vs control and 5-HT vs control comparisons. In G and H, black arrows: interactions unchanged with treatment, cyan arrows: interaction lost with treatment, magenta arrows: interaction gained/emerged with the treatment. See also Figure S3.

We found that *bdnf* was predominantly expressed in neuronal clusters (90% in neurons and 8% in glial cells; Figure3E; FigureS3H), while its receptor *ntrk2* (Abbate et al., 2014; Lucini et al., 2018) was expressed mainly in neurons (68% neuronal and 28% glia; Figure3E; FigureS3H), while another BDNF receptor *ngfra* (Abbate et al., 2014; Cacialli et al., 2016; Lucini et al., 2018) was mostly glial (93% in glia and 5% in neurons; Figure3E; FigureS3H). Therefore, we hypothesized that 5-HT dependent regulation of *bdnf* expression might signal to the glial cells through *ntrk2* and *ngfra* in adult zebrafish brain.

To further investigate if BDNF signaling through Ntrk2 or Ngfra would affect NSCs, we used an in silico interaction map analyses we recently developed (Cosacak et al., 2019). According to this analyses, if BDNF would have a potential interaction between neuronal and glial (progenitor) clusters, we would see an in silico interaction between these cells. Alternatively, if *ntrk2* or *ngfra* could constitute a crosstalk between neuronal and glial clusters, we would be able to see an interaction (see (Cosacak et al., 2019) for details of interaction mapping). For interaction analyses, we used the spatial organization map of adult zebrafish NSCs in the telencephalon (Figure 3F), distinguished the cell types based on our previous findings ((Cosacak et al., 2019)) by using machine learning algorithms, and compared the three treatments with controls (IL4 vs. control, Amyloid vs. control, and 5-HT vs. control) (Figure 3G,H; Figure S4A). For mapping, we used the following three criteria: (1) we took either the *bdnf* as a starting point (i.e.: the cells expressing *bdnf* are matched with the cells expressing any *bdnf* receptor, or the *bdnf* receptors *ntrk2* and *ngfra* were taken as starting points (i.e.: only *ntrk2* or *ngfra*-expressing cells were matched with cells expressing any ligand for those receptors (Figure3H for *ngfra* and Figure S4A for *ntrk2*)); (2) charted the potential interactions as arrows starting from *bdnf*-expressing clusters to *bdnf* receptor-expressing clusters (Figure 3G) or to *bdnf* receptor-expressing clusters from clusters expressing their ligands (Figure3H), and (3) depending on the change of the interaction after any treatment, we color-coded the interactions (black arrows indicate interactions that are unchanged by the treatment, cyan arrows indicate the interactions that are lost after treatment, and magenta arrows indicate interactions that emerged after a particular treatment). After these analyses, we found that especially two clusters of progenitor cells (PC0 and PC2, which are located to dorsal and medial region of the telencephalon (Figure 3F) and (Cosacak et al., 2019)) were among the clusters that had the highest number of arrows pointing towards them (Figure3G). Interestingly, the majority of the potential interactions were newly generated after Amyloid or IL4 treatment while those interactions are mainly lost after 5-HT treatment (Figure 3G, green circles), suggesting that the regulation of progenitor cells (neural stem cells) by *bdnf* could be promoted by Amyloid or IL4 treatment and suppressed by 5-HT, which is consistent with our hypothesis. Additionally, to support our findings, we performed independent mapping analyses of *bdnf* receptors *ntrk2* and *ngfra*. We constructed interaction maps for these receptors and found that *ngfra* might be the main receptor for *bdnf* signaling in adult zebrafish telencephalon as two NSC clusters (PC0 and PC2) were at the center of the interaction maps with many potential interactions (Figure 3H) while *ntrk2* interaction map provided only small number of interactions to another glial cluster (PC7) (Figure S4A). We observed that IL4 activates many interaction routes to NSC clusters while 5-HT almost diminishes all the interactions between NSCs and other cells (Figure 3H). These in silico analyses suggest that IL4 promotes the interactions between neurons and NSCs while 5-HT rather suppresses such interactions.

To verify our in silico analyses for interaction mapping on single cell transcriptomics data and to investigate the changes in the expression of *bdnf*, we performed in situ hybridization on control (Figure 4A), 5-HT-injected (Figure 4B) and IL4-injected brains (Figure 4C). The expression of *bdnf* in control brains was in periventricular zone (PVZ) proximal to the NSCs, which confirmed our single cell transcriptomics data. The expression of *bdnf* was almost abolished by 5-HT injection (Figure 4B) while it is enhanced by IL4 (Figure 4C), suggesting that 5-HT suppresses NSC plasticity through reducing the *bdnf* signaling.

**Figure4:**
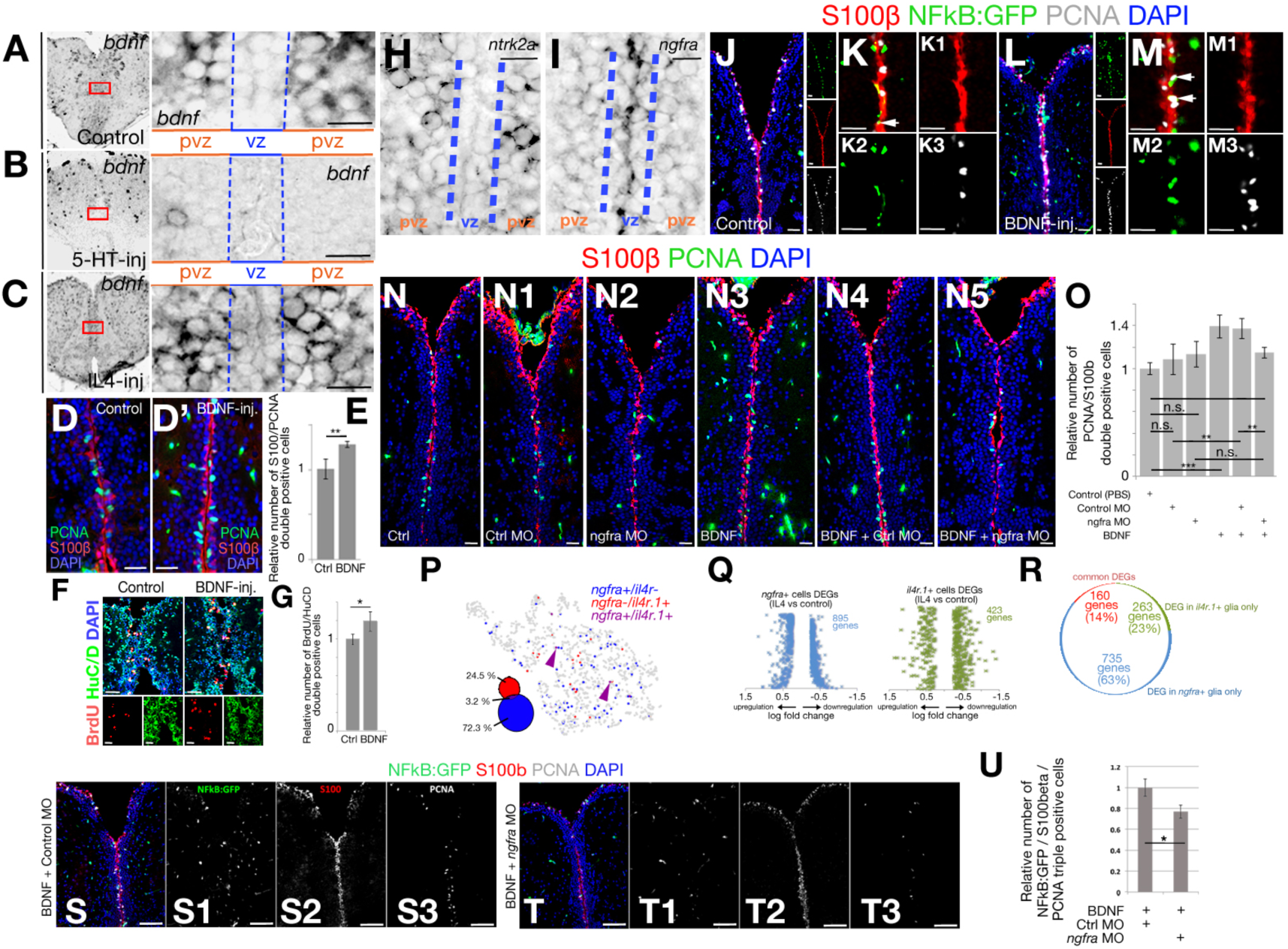
Serotonin regulates periventricular *bdnf* expression, which regulates neural stem cell plasticity through *ngfra.* (A-C) In situ hybridization (ISH) for *bdnf* in control (A), 5-HT-injected (B) and IL4-injected (C) brains. Red rectangles are enlarged to the right of the main panels. vz: ventricular zone, pvz: periventricular zone. Note significant reduction of *bdnf* expression after 5-HT. (D, D’) IHC for PCNA and S100β in control (D) and BDNF-injected (D’) brains. (E) Quantification of proliferating glial cells after BDNF injection. (F) IHC for BrdU and HuC/D in control and BDNF-injected brains. (G) Quantification of newborn neurons at 14dpi after BrdU treatment at Day2 and Day3 after PBS and BDNF injection. (H) ISH for *ntrk2*, which is expressed in periventricular neurons (pvz) but not in neural stem cells in the ventricular zone (vz). (I) ISH for *ngfra*, which is expressed in vz. (J) IHC for S100β, NFkB-driven GFP, and PCNA in control brains. To the right of the larger panel are single fluorescence channels. (K) High-magnification of the medial region of J without DAPI. (K1-K3) Single fluorescent channels of K. (L) IHC for S100β, NFkB-driven GFP, and PCNA in BDNF-injected brains. To the right of the larger panel are single fluorescence channels. (M) High-magnification of the medial region of L without DAPI. (M1-M3) Single fluorescent channels of M. (N-N5) IHC for S100β and PCNA in control (N), control morpholino-injected (N1), *ngfra* morpholino-injected (N2), BDNF-injected (N3), BDNF + control morpholino-injected (N4) and BDNF + *ngfra* morpholino-injected (N5) brains. (O) Quantification for the relative number of proliferating glial cells. (P) Co-representation of *ngfra* and *ilr4* expressions on glial tSNE plot. (Q) Differentially expressed gene (DEG) plots for *ngfra-*positive and *il4r*-positive neural stem cells after IL4 treatment. (R) Pie-chart distribution of unique differentially expressed genes. Note that the overlapping genes constitute only 14% of all DEGs. (S) Immunohistochemistry for S100b, NFkB-driven GFP and PCNA in BDNF + control morpholino-injected (S-S3), and BDNF + *ngfra* morpholino-injected brains (T-T3).(U) Quantification graph for relative numbers of proliferating stem cells with active NFkB signaling. Scale bars equal 100 µM. Data are represented as mean ± SEM. See also Figure S4-S8.

Given that 5-HT suppresses *bdnf* expression and NSC proliferation while IL4 enhances *bdnf* expression and NSC proliferation, we hypothesized that *bdnf* would also enhance NSC plasticity by increasing cell proliferation. To test this, we injected BDNF into control zebrafish brains (Figure 4D-E) and observed that BDNF indeed increases the number of proliferating NSCs (Figure 4E). The enhanced proliferation of NSCs by BDNF is also translated into increased neurogenesis as BDNF-injected brains produce more neurons compared to control-injected zebrafish brains (Figure4F, G), indicating that BDNF enhances NSC plasticity in adult zebrafish brain.

If BDNF would act directly on NSCs, its receptor must have been present in the target cells. Therefore, we performed in situ hybridization for *ntrk2* and *ngfra* in adult zebrafish brains. We observed that *ntrk2* mRNA is expressed mainly in periventricular region of the telencephalon (Figure 4H and Figure S4B), which is supported by immunohistochemical staining for NTRK2 protein (Figure S4C). On the contrary, *ngfra* is expressed mainly in ventricular zone where NSCs reside (Figure 4I, Figure S4). These findings are perfectly matching with our single cell transcriptomics data (Figure 3E, Figure S3H), and suggest that *ngfra* is the primary receptor for *bdnf* in NSCs and *ntrk2* is the primary receptor in neurons. To test this hypothesis, we injected BDNF into adult zebrafish brain and determined the downstream effectors of NTRK2 and NGFRA signaling pathways. To determine the cells responding to BDNF/NTRK2 signaling, we detected the downstream effector pAkt (Huang and Reichardt, 2003) and observed that after BDNF injection, pAkt is almost exclusively present in periventricular cells and parenchymal neurons (few speckles in ventricular region constitute less than 0.1% of the pAkt signal when compared to the intensity and number of cells outside the ventricular region) (Figure S4D,E). On the contrary, compared to control-injected brains, BDNF injection increased the activity of the NFkB reporter (Kanther et al., 2011), which is the downstream effector of NGFR signaling (Hamanoue et al., 1999) in the ventricular region where NSCs reside (Figure 4J-M). Additionally, consistent with the results of BDNF injections, Aβ42 or IL4 injections increased but 5-HT injection reduced the NFkB signaling and the number of NFkB-positive proliferating NSCs (S100b/PCNA/NFkB:GFP-triple positive cells; Figure S4G,H). These results indicate that NGFR-mediated intracellular signaling is the primary route for the 5-HT-dependent BDNF activity on NSCs in adult zebrafish brain.

If our hypothesis were true and BDNF would regulate NSC proliferation through NGFRA and NFkB signaling, knocking-down *ngfra* receptor after BDNF injection would suppress the increase in NSC proliferation by BDNF as well as the increase in NFkB signaling. To test this hypothesis, we knocked down *ngfra* by using morpholino oligonucleotides and determined the extent of proliferating NSCs (Figure 4N-N5). Compared to control brains, BDNF increased the NSC proliferation; control morpholino did not alter this increase while *ngfra* morpholino diminished the increased NSC proliferation upon BDNF injection (Figure4O).

Since BDNF/NGFRA signaling is affected by IL4 that also acts directly on NSCs through IL4R (Bhattarai et al., 2016), we hypothesized that the effect of IL4 on NSCs could be mediated through IL4R (directly by IL4/IL4R interaction) and NGFR (through 5-HT/BDNF/NGFRA/NFkB axis) distinctly, and IL4R-positive and NGFRA-positive glia would constitute two functional subtypes of NSCs. To determine whether IL4R-positive and NGFRA-positive NSCs are distinct subtypes of NSCs in adult zebrafish brain, we plotted both cell populations on the same tSNE plot (Figure 4P). We observed that only 3,2% of *ngfra-*positive NSCs were also *il4r1-*positive (24,5% are only *il4r*-positive and 72.3% are only *ngfra*-positive), indicating that these two populations are likely to represent two functional subtypes of NSCs. We further hypothesized that if *ngfra*-positive and *il4r*-positive NSCs would constitute different subtypes, their response to particular treatments would also lead to distinct differential gene expression profiles. To address this question, we determined the overlap between differentially expressed genes in *il4r*-positive and *ngfra*-positive NSCs (Figure 4Q). We determined that after IL4 treatment 423 and 895 genes are differentially expressed in *il4r*+ (Dataset S5) and *ngfra*+ (Dataset S6) NSCs, respectively (Figure 4Q). Only 14% of the differentially expressed genes are common in these cell populations while the rest are differentially expressed in only one of the cell types (Figure 4R). Similarly, *ngfra* knockdown reduced the number of proliferating NSCs where NFkB signaling is active (Figure 4S-T3). These results indicate that neuron-glia interaction through BDNF regulates NSC proliferation and neurogenesis through NGFRA/NFkB signaling (Figure 5).

**Figure 5:**
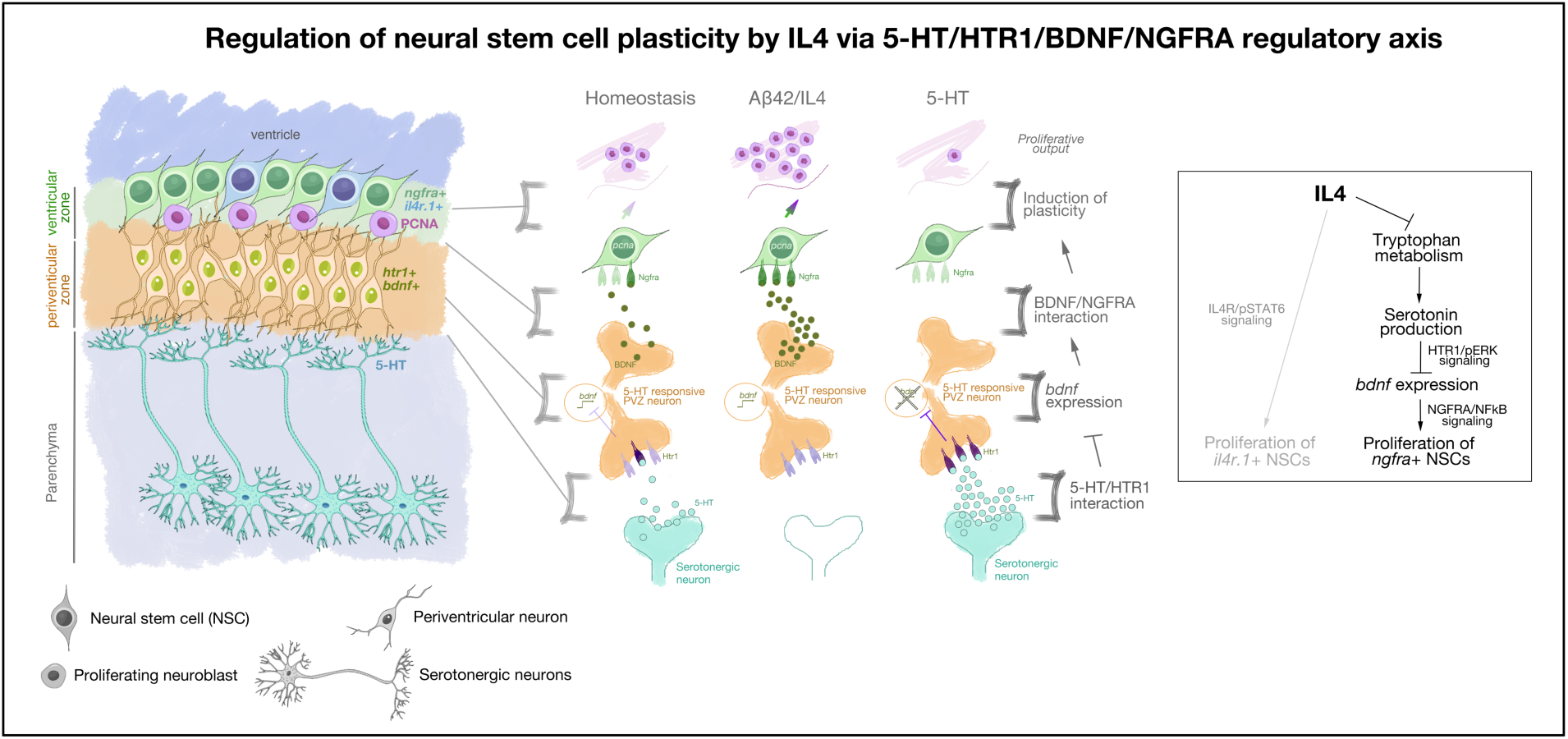
Schematic view of the findings on how neuron-glia crosstalk regulates Alzheimer’s-induced neurogenesis in adult zebrafish brain.

To explore the evolutionary conservation of our findings in healthy and Alzheimer’s disease conditions, we determined the expression of BDNF, NTRK2 and p75/NTR (NGFRA ortholog in mouse) in wild type mouse and APP/PS1dE9 AD model (Figures S5-7). Compared to 12 month-old control mouse brains, age-matched APP/PS1dE9 mouse displayed reduced SOX2 and increased GFAP (Figures S5-7) that is indicative of reduced neurogenic ability and increased gliosis. We found that in mouse cortex and dentate gyrus (DG), BDNF is mainly expressed by non-glial cells (Figure S5C-D’; Figure S6A-F), which is supported by the previous studies (Habib et al., 2016; Tabula Muris et al., 2018) and this pattern is not altered in AD brains (Figure S5E-F’; Figure S6G-L).

NTRK2 is expressed in the cortex and DG, again mainly in non-glial cells but few GFAP-positive astrocytes were NTRK2-positive in both the wild type and AD mouse brains with no clear change in the expression pattern between healthy and diseased brains (Figure S5G-J’). Overall, NTRK2 is expressed mainly by neurons, but also few astrocytes and microglia (FigureS6M-P). We found that p75/NTR is mainly expressed in neurons in the cortex and the DG, but NSC niche in the DG (subgranular zone, SGZ) does not express p75/NTR (Figure S5K-N’; Figure S7). In both wild type and APP/PS1dE9 mouse, we could detect BDNF, NTRK2 and p75 expression; however, increased number of GFAP-positive astrocytes AD brains did not correlate with the expression of these proteins, suggesting that BDNF signaling might not regulate NSC proliferation in mouse brains.

To test this hypothesis, we investigated the effect of BDNF in wild-type and AD mouse brains by injecting BDNF into the mouse brains (Figure S5O-X; Fig S8). BDNF injection was performed in one hemisphere of the mouse brain, whereas the other hemisphere was used as a control with PBS injection. We found that BDNF injection increased the overall proliferation levels in the brain as compared to PBS-injected hemispheres in both wild type and APP/PS1dE9 brains (Figure S5O-V2). In addition we also checked the cell types that increase their proliferation levels.

To identify the BDNF-responsive proliferating cells, we performed co-staining with astrocyte marker GFAP and found that Ki67+ cells are GFAP negative, suggesting that BDNF responsive proliferating cells are not astrocytes (Figure S5O-R2; Figure S5W; Figure S8A-C), Next, we performed Ki67 co-staining with microglial marker Iba1 and found that almost all Ki67-positive cells are Iba1-positive, suggesting that BDNF induces microglial proliferation resulting in microgliosis but not NSC proliferation (Figure S5S-V2; Figure S5X; Figure S8D-F). Consistent with this hypothesis, we also found that SOX2-positive cells did not alter their proliferation levels after BDNF (data not shown), suggesting that BDNF does not alter NSC proliferation in mouse brains in healthy and AD conditions. This finding is consistent with the previous reports (Choi et al., 2018; Galvao et al., 2008; Reumers et al., 2008). Additionally, due to the lack of p75NTR expression in NSCs present in SGZ of the mouse hippocampus, BDNF/p75NTR signaling, which enhances proliferative output of NSCs and neurogenesis in zebrafish brain, is not an active signaling mechanism in mouse brains. With these results, we propose that zebrafish utilizes special circuit mechanism that uses serotonin-BDNF signaling to enhance the NSC plasticity and to induce neurogenesis through neuronal intermediates; however, BDNF signaling is not regulating NSC plasticity and neurogenesis in mammalian brains while its effect is mainly on the neuronal survival and regulation of immune reaction (Chan et al., 2008; Galvao et al., 2008; Hamanoue et al., 1999; Huang and Reichardt, 2003; Takach et al., 2015; Yeiser et al., 2004).

## Discussion

Our results identify a previously uncharacterized regulatory circuit mechanism that involves neuronal intermediates for NSC plasticity of zebrafish brain in Alzheimer’s disease (AD) conditions. This mechanism involves the suppressive effect of Serotonin on periventricular neurons that express *bdnf*, which directly regulates a subset of NSCs expressing BDNF receptor *ngfra*. Amyloid-induced Interleukin-4 suppresses the initial serotonergic output and therefore potentiates the *bdnf* expression and stem cell proliferation. We propose that induced expression of IL4 generates a disease-associated NSC niche environment that is supportive of NSC proliferation.

Serotonin affects neurogenesis, stem cell proliferation and normal functioning of neural circuitry; however, its effects on NSCs and neurogenesis are controversial (Alenina and Klempin, 2015). In zebrafish, Serotonin positively affects NSC proliferation in midbrain but not the hypothalamus (Perez et al., 2013), and promotes spinal motor neuron regeneration (Barreiro-Iglesias et al., 2015). Although depletion of serotonin reduces adult neurogenic outcome in rats (Brezun and Daszuta, 1999), elevated levels of serotonin in serotonin transporter (SERT) deficient mice did not affect NSC proliferation and neurogenesis (Bengel et al., 1998; Schmitt et al., 2007). In mesencephalic and hippocampal progenitors of mouse brain, inhibition of serotonin receptors or knocking out serotonin-producing enzymes to generate hypo-serotonergic phenotypes had positive effects on neurogenesis (Diaz et al., 2013; Klempin et al., 2013; Parga et al., 2007; Sachs et al., 2013) ascertaining a negative role for serotonin on NSC plasticity. These findings indicate that the effects of serotonin on NSCs are context-dependent and may run through intermediary secondary signaling mechanisms, which supports our findings. Additionally, in majority of the studies, the specific receptors for serotonin were not investigated in NSCs; therefore our study provides a detailed delineation of the signaling cascades that link serotonergic input to the regulation of stem cell proliferation via intermediates, in our study by BDNF.

The interaction between serotonin and BDNF is partially understood. In stress-related disorders depletion of serotonin reduces brain BDNF levels (Chen et al., 2007; Yoshida et al., 2012; Zhou et al., 2008). On the contrary, hypo-serotonergic mouse (*Tph2*−/− or *Pet1*−/−) showed increased BDNF levels in hippocampus and prefrontal cortex (Diaz et al., 2013; Klempin et al., 2013; Kronenberg et al., 2016; Sachs et al., 2013) and humans with serotonin transporter deficiency (hyper-serotonin) reduced the availability of BDNF (Molteni et al., 2010), which enhances NSCs proliferation and neurogenesis (Arsenijevic and Weiss, 1998; Numakawa et al., 2017). Our findings support the negative role of serotonin on BDNF signaling and NSC plasticity.

The effects of BDNF in AD are mainly on neuronal survival rather than neurogenesis or NSC plasticity. Although BDNF expression correlates with increased neuronal survival (Lee et al., 2002; Quesseveur et al., 2013; Waterhouse et al., 2012), and may have positive impact on increased net hippocampal neurogenesis and better cognitive functioning (Rossi et al., 2006; Wrann et al., 2013), several studies showed that these effects are not directly related to enhanced neurogenesis (Galvao et al., 2008; Reumers et al., 2008). In fact, in a recent study, in 5X-FAD AD model of mice, increasing adult neurogenesis and simultaneous BDNF treatment increases cognitive output yet adult neurogenesis or BDNF alone are not sufficient to do so (Choi et al., 2018), suggesting that a BDNF-dependent neuronal survival cascade is required to counteract the symptoms of AD after an independent stimulation of neurogenesis. In zebrafish, in contrast to mammals, BDNF directly regulates NSC plasticity and neurogenesis. Our results in APP/PS1dE9 mice also supported the findings in mice (Choi et al., 2018) and indicate that zebrafish has an intrinsic and natural ability to kick-start NSC proliferation and neurogenesis, which could teach us how to modulate mammalian brains to initiate “regeneration” of lost neurons in AD.

BDNF expression is abundant in mammalian brains (Leibrock et al., 1989) and its main receptor is *TrkB/Ntrk2*, which is predominantly expressed in neurons (Klein et al., 1991) in the cortex and DG as well as in few astrocytes and microglia as determined by single-cell sequencing (Habib et al., 2016; Tabula Muris et al., 2018). BDNF also has binding affinity towards p75 receptor, which is primarily expressed by neurons in mammals (Habib et al., 2016; Tabula Muris et al., 2018). Yet, in zebrafish, *ngfra* was predominantly expressed in NSCs and were responsible for NSC plasticity. Zebrafish glial cells responded to BDNF via *ngfra* receptor, and activated proliferation and neurogenesis; whereas, mouse glial cells responded to BDNF by reactive astrogliosis and microgliosis. Therefore, our results also propose that the BDNF activity could be specialized to distinct receptor signaling pathways in vertebrate brains. For instance, BDNF-NGFRA signaling could be a factor underlying the regenerative and plastic nature of the NSCs in regenerating organisms such as zebrafish but not in non-regenerating organisms like mammals, where BDNF mostly acts through TrkB/Ntrk2 signaling. Further studies of this regulatory mechanism in other regenerating organisms would be instrumental in testing our hypothesis. Furthermore, inducing expression of *ngfra* in stem cell populations in mammalian brains could be a potential way to impose neurogenic plasticity to astrocytes in AD conditions.

We found that IL4 suppresses the serotonin production in neurons that do not express *il4r*, suggesting that IL4 regulates serotonin availability indirectly. Indeed, pro-inflammatory cytokines such as IL-1β, TNFα, IFNα and IFNγ can increase 5-HT levels and the expression of 5-HT transporter SERT, which positively correlates with 5-HT levels in the brain (Baganz and Blakely, 2013; Linthorst et al., 1996; Masson and Hamon, 2009; Masson et al., 1999; Mossner et al., 1998; Ramamoorthy et al., 1995; Shintani et al., 1993). Theoretically, anti inflammatory factors would be expected to reduce 5-HT levels in the brain due to their role in suppression of pro-inflammatory factor expression. Indeed, IL4 reduces the 5-HT or SERT levels (Mossner et al., 2001; Pannell et al., 2014; Park et al., 2015; Wachholz et al., 2017), supporting our findings. Therefore, in adult zebrafish brain, AD conditions elicit an inflammatory regulation by IL4 on serotonin, which translates into a neurotransmitter response in a specific neuronal subtype that directly regulates a subpopulation of NSCs.

Finally, we identified two functionally distinct populations of NSCs that are both responsive to IL4 but through different receptor signaling (IL4R/STAT6 and BDNF/NGFRA) and through distinct gene regulation (Figure3X,Y). This finding suggests that NSC heterogeneity is an important determinant of how a certain type of neuropathology and a particular signaling pathway would affect certain subtypes of cells differentially (in AD regulation of proliferation by IL4 in two distinct branches of regulation (Figure5)). This understanding would be instrumental in designing targeted therapy options for AD in humans and may lead to more detailed analyses of NSC subtypes in experimental mammalian models of AD. Additionally, the role of Interleukin-4 and Serotonin in regulating diverse signaling mechanisms in NSCs opens up interesting research avenues as to whether modulation of neuroinflammatory milieu in mammalian brains would kick-start a “regeneration” response by mobilizing the endogenous reservoir of NSCs, which is a controversial subject (Arellano et al., 2018; Cipriani et al., 2018; Rodriguez and Verkhratsky, 2011; Sorrells et al., 2018; Tincer et al., 2016; Waldau and Shetty, 2008; Ziabreva et al., 2006). We propose that zebrafish can be used to address neurogenesis-related questions in disease conditions and could serve as a useful experimental model for investigating the molecular mechanisms of neural stem cell plasticity.

## Acknowledgements

This work was supported by German Center for Neurodegenerative Diseases (DZNE) and Helmholtz Association through Young Investigator Award (VH-NG-1021, C.K.) and Deutsche Forschungsgemeinschaft (DFG) (KI1524/6, KI1524/10, and KI1524/11, C.K.). We would like to thank to John F. Rawls and Steven Renshaw for NFkB:GFP reporter zebrafish line, Tomohisa Toda and Gerd Kempermann for critical comments on the manuscript, and to Andreas Dahl, Susanne Reinhardt, Andreas Petzold for single cell sequencing, and to Klaus Fabel and Gerd Kempermann for providing APP/PS1dE9 mouse.

## Author contributions

C.K. and P.B. conceived and designed the experiments. P.B. performed experimental procedures for injections, immunohistochemical stainings, quantifications and imaging in zebrafish. V.M. performed injections into the mouse brains and sectioning. P.B. performed mouse immunohistochemical stainings, quantifications and imaging. K.B., and S.D.P helped in situ hybridizations. SY helped immunohistochemical stainings and quantifications. M.I.C. performed bioinformatics analyses including the whole transcriptome and single cell sequencing analyses. Y.Z. provided the Amyloid peptide, K.B. and N.G. performed histological sectioning of the APP/PS1dE9 mouse. C.K. acquired funding, supervised the study, C.K. and P.B. wrote the manuscript, prepared the figures, and edited the manuscript.

## Experimental Procedures

### Animal Experiments

#### Ethics statement

Animal experiments were carried out in accordance with the relevant permits from the Landesdirektion Sachsen, Germany (TVV-52/2015 and TVV-87/2016).

#### Animal Handling and Husbandry

For zebrafish studies, 6-8 months old WT-AB strain, Tg(her4.1:GFP) and Tg(NFkB:GFP) fish of both genders were used. For each set of experiments, same clutch of fish were randomly distributed for different experimental groups. Fish were maintained at 14 hour light / 10 hour dark cycle at constant 28 °C tanks. All animal experimentation was performed according to the official guidelines. For mouse studies, APP/PS1 mice was kept and bred according to established protocols. 12-months-old transgenic mice and WT littermates (males only) were used for the experiments. These mice were maintained following the institutional guidelines in standard animal housing conditions.

### Peptide synthesis, cerebroventricular microinjection in adult zebrafish

Amyloid-β42 (Aβ42) synthesis was performed as described (Bhattarai et al., 2016). Cerebroventricular microinjection in adult zebrafish brain were performed as previously described (Bhattarai et al., 2017a; Bhattarai et al., 2016; Bhattarai et al., 2017b). PBS (1μl), Aβ42 (1 μl, 20 μM), IL4 (1 μl, 10 μg/ml), BDNF (1 μl, 100ng/ml), 5-HT (1 μl, 100 μM), ngfra morpholino (1μl, 10uM) were injected. See Table S1 for further information.

### BrdU treatment

Experimental zebrafish were immersed in freshly prepared 10 mM BrdU (Sigma-Aldrich) solution in E3 for 8 hours per day at 2 and 3 days post-injection (dpi). Fish were sacrificed at 14 dpi, and heads were subjected to histological preparations.

### Zebrafish tissue preparation, immunohistochemistry and in situ hybridization

Zebrafish tissue preparations, cryosectioning, mRNA probe synthesis and stainings were performed as previously described (Bhattarai et al., 2016; Kizil et al., 2012a; Kizil et al., 2012b). See Table S1 for further information on reagents.

### BDNF Injection into adult mouse brain

For mouse experiment, 100 ng of BDNF (1 μl, 100μg/ml) was injected into one hemisphere of the adult mouse brain, and the other hemisphere was injected with 1 µl PBS as a control. The injection procedure was carried out according to previously established protocol (Artegiani et al., 2011). The mouse was placed in an induction chamber and anesthetized using a mix of oxygen and isoflurane flow. After the animal was recumbent, the gas flow directed to the nosecone of the stereotaxic frame. The mouse was then rapidly positioned on the nosecone to have constant supply of isoflurane and oxygen. The ear bars were fixed to immobilize the head. An analgesic was subcutaneously injected prior to surgery to minimize any possible pain after the recovery from anesthesia. During the entire surgery animal was placed on a pre-warmed heat-pad to prevent hypothermia. To prevent dehydration of cornea and further blinding of mouse, eyes wer covered with a protective ointment. All tools used during the surgery were sterilized using ethanol and/or Microzide. Using bregma as a reference point, the coordinates for the injections into the Dentate Gyrus were identified on both sides of the brain and holes were carefully drilled in the skull. The capillary was then slowly inserted until 1900 um of depth from skull level and 1 µl of BDNF (100μg/ml) was injected at 200 nl/min speed. After injection, the capillary was kept inside for 5 minutes to prevent backsplash and then slowly retracted. The same procedure was performed on the other hemisphere to inject 1 µl of PBS. The ear bars were then released, the skin was wetted with PBS to regain the elasticity and stitched. The wound was then treated with a disinfecting solution (Povidone). Mouse was placed in an individual cage under a red light with water and pre-soaked food (to make it easier for animal to bite after the surgery). Another injection of analgesic was performed 24 hours post-surgery. Mouse were sacrificed at 48 hours after injection and subjected for brain collection and further processing.

### Mouse tissue preparations and stainings

For brain collection the adult mice were anesthetized with an intraperitoneal injection of a mixture of Ketamine and Xylazine (0,25 mL per 25g of body weight) and then intracardiacally perfused with NaCl 0.9% followed by cold freshly prepared 4% PFA (in phosphate buffer, pH 7.4). After perfusion brains were harvested and fixed with 4% PFA for 24 hours at 4°C and then washed 3 times with PBS. For cryopreservation of tissue fixed brais were further incubated overnight in 30% Sucrose solution in PBS at 4°C. Free-floating sections of 40µm thickness were prepared using microtome (Leica SM2010R) and were collected in 6 consecutive series in a Cryopreservation Solution (CPS) in a 24-well plate. Sections were then stored at −20 °C for further experiments. Prior to immunohistochemistry the free-floating sections were washed in PBS 3 times, blocked in 10% Donkey or Goat Serum with 0.2% TritonX for 1 hour at room temperature and incubated overnight at 4 °C with the desired primary antibody of defined dilution in 3% Donkey or Goat Serum with 0.2% TritonX. Sections were washed for 3 times within 1 hour and incubated for another 4 hours at room temperature with the correct secondary antibody coupled to a desired fluorophore. After short wash samples are then incubated in DAPI diluted in PBS (1:5000) for 10 minutes. Another 3 washing steps were done and samples were then mounted on the charged glass slides (Thermo Fisher, SuperFrostTM). After mounting slides are left to dry and covered with a coverslip using Aqua Mount. See Table S1 for further information on reagents.

### Imaging and quantifications

Images were acquired using ZEN software on a Zeiss fluorescent microscope with ApoTome or Zeiss AxioImager Z1 and analyzed using ZEN or ImageJ software. Quantifications were performed in double blind fashion. Stereological quantifications in zebrafish were performed by manual counting on one third of the whole sections from the Telencephalon (caudal olfactory bulb until the rostral optic tectum). Quantifications for serotonergic densities were performed using a 3D object counter module of ImageJ software. At least 4 animals with minimum 8 histological sections per animal were used for stereological analyses in all in situ hybridization and immunohistochemical stainings unless otherwise stated.

### Statistical analyses

All experiments were performed at least with two replicates. For comparison of two particular experimental groups, a two-tailed student’s t-test was performed. For comparison of multiple experimental groups, one-way ANOVA with Tukey’s post-test were performed. Statistical analyses were performed using Excel software and GraphPad Prism. p-values less than 0.05 were considered significant. Error bars in Fig.s indicate the standard error of means. Significance indicated by ∗ (p < 0.05), ∗∗ (p < 0.01), and ∗∗∗ (p < 0.001), n.s. (not significant, p > 0.05). No sample set was excluded from the analyses unless the histological sections were damaged severely during the acquisition of the sections (constitutes less than 4% of all sections analyzed).

### Cell dissociation, sorting, whole transcriptome and single-cell sequencing

The telencephalon was dissected using ice-cold PBS and Neural Tissue Dissociation Kit (Miltenyi) as described previously (Bhattarai et al., 2016). Deep sequencing for whole transcriptome was performed as described (Bhattarai et al., 2016; Papadimitriou et al., 2018). For single cell sequencing, isolated cells were passed through a 40-µm cell strainer. Viability indicator (Propidium iodide) and GFP were used to sort live cells. Cell suspension was loaded into the 10X Chromium system (Zheng et al., 2016). 10X libraries were prepared as per the manufacturer’s instructions. The raw sequencing data was processed by the cell ranger software provided by the 10X genomics with the default options. The reads were aligned to zebrafish reference transcriptome (ENSEMBL Zv10, release 91). Analyses matrices were used as input for downstream data analysis by Seurat (Butler et al., 2018). Analyses were performed as described (Cosacak et al., 2019).

### Data Analysis by Seurat

Read10X function was used for generated matrices. Cells that contain more than 1000 UMI and more than 200 unique genes were taken into consideration. The data was normalized by using “LogNormalize” method, and was scaled with “scale.factor = 1e4”. Variable genes were detected with FindVariableGenes using the default options. The top1000 most variable genes from each sample were merged, and CCA analysis was performed on the variable genes. A merged Seurat object was created with RunMultiCCA function, using num.ccs = 50. The canonical correlation strength was calculated using num.dims = 1:20. The reduction was performed by reduction.type = “pca”, dims.use = 1:20, then the data subset by using “var.ratio.pca” and “accept.low = 0.5”. The samples were aligned using dims.align = 1:20. The cell clusters were found using aligned CCA and 1:20 dims, with higher resolution 2.5. Clusters were shown on 2D using t-SNE function. We used the genes from literature (Dataset S4) and markers found by Seurat functions to define our cell types. For that, we generated FeaturePlot for all marker genes and checked the expression and specificity of the marker to define cell types.

### Pathway analyses and interaction mapping by machine learning

We used all marker genes with False Detection Rate < 0.1 for Gene Ontology analysis and KEGG pathway analysis using GOStats (1.7.4) (Falcon and Gentleman, 2007) and GSEABase (1.40.1), p-value < 0.05 as thresholds. To determine the differentially expressed genes (DEGs), we used FindMarkers function using cell cluster that have at least 3 cells from all samples. Then, we used the p-value <0.05 for significantly expressed genes. These genes were used for GO and pathway analysis using the scripts that are available on https://kizillab.org/resources.

Cell types in Serotonin single-cell sequencing dataset were identified by using machine learning algorithms. Top-1000 most variable genes for three samples (PBS, AB42 and IL4 as previously described (Cosacak et al., 2019)) were determined. These genes were used to train RandomForest algorithm to learn cell types from existing samples. The cells in 5-HT treated samples were predicted accordingly. We identified marker genes, differentially expressed genes (DEGs), interaction maps as described before (Cosacak et al., 2019).

To identify DEGs in specific cell types, we added new “idents” to the R-object as the following: Treatment (PBS, AB42, IL4 or 5HT)_Main Cell Types (Im, PC, NN, OPC/OD1 or OPC/OD_2)_receptor name. For instance; AB42_PC_htr1a was compared to PBS_PC_htr1a cells. The DEGs in each of these cells were identified after each treatment using FindMarkers function in Seurat. p-value < 0.05 and |avg_logFC| >= 0.25 were used for significantly expressed genes, p-value < 0.05 was used for over-represented GO terms and KEGG pathways as described (Cosacak et al., 2019).

### Data availability

GEO Accession numbers for sequencing datasets are as follows: single cell sequencing: GSM3334110 (control), GSM3333461 (Amyloid-beta42), GSM3333764 (Interleukin-4) (as described in (Cosacak et al., 2019)) and GSM3334111 (5-HT/Serotonin); whole transcriptome deep sequencing dataset: GSE124162. All R-Codes are available on kizillab.org/resources.

## Supplementary Materials

**Figure S1:**
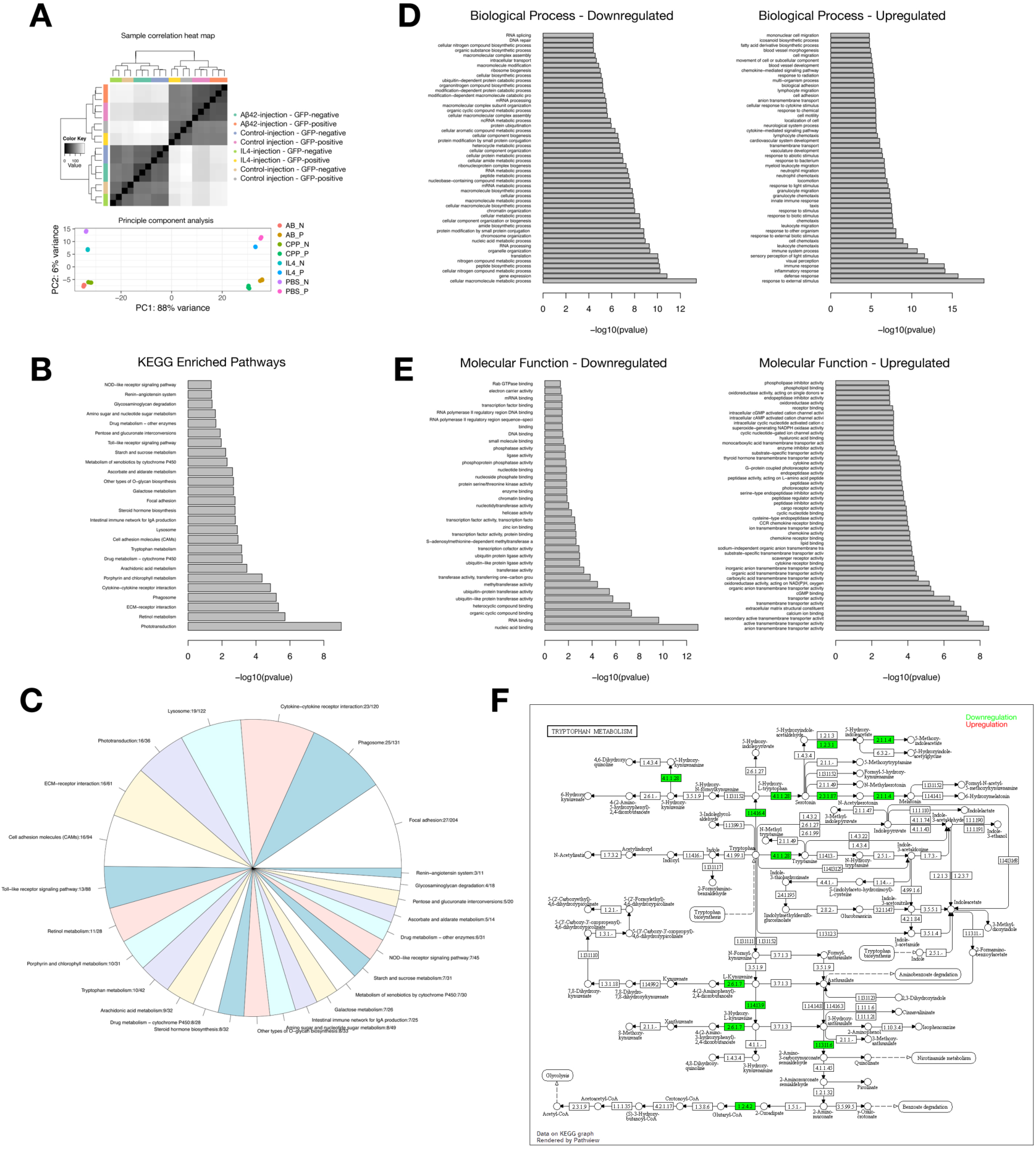
Analyses of whole transcriptome sequencing after IL4 treatment. (A) Sample correlation heat map and principle component analyses. (B,C) KEGG enriched pathways as list (B) and pie chart (C). (D) GO terms for biological process. (E) GO terms for molecular function. (F) Tryptophan metabolism in detail. Green: downregulated enzymes, red: upregulated enzymes after IL4. Related to Figure 1.

**FigureS2:**
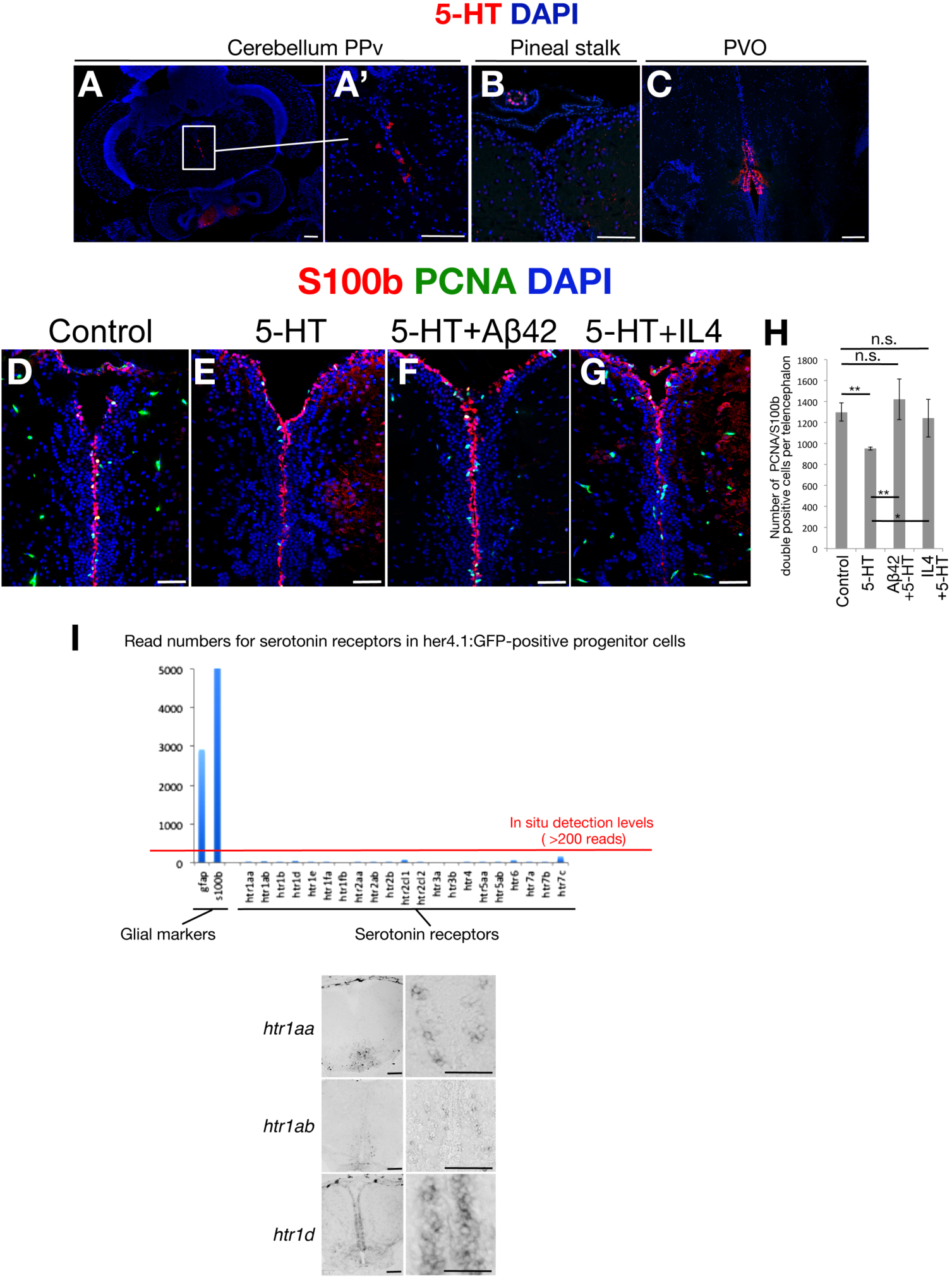
Amyloid-beta42 and IL4 antagonize the effect of 5-HT on neural stem cell plasticity. (A-C) 5-HT immunostaining in caudal regions of the adult zebrafish brain: Cerebellum (A, A’), pineal stalk (B), and caudal raphe, PVO (C). (D-G) Immunohistochemistry for S100beta and PCNA on control (D), 5-HT-injected (E), 5-HT + Ab42-injected (F) and 5-HT + IL4-injected (G) zebrafish brains. (H) Quantification of proliferating glial cells in all conditions. (I) Read numbers of all serotonin receptors in her4.1-positive cells (progenitor cells) in the adult zebrafish telencephalon as a graphical representation that is derived from deep sequencing results. Note that progenitor cells/glia do not express serotonin receptors as shown in ISH panels of *htr1aa, htr1ab and htr1d*. Glial markers *gfap* and *s100* are given as positive controls. Scale bars equal 100 µM. Related to Figure 2.

**FigureS3:**
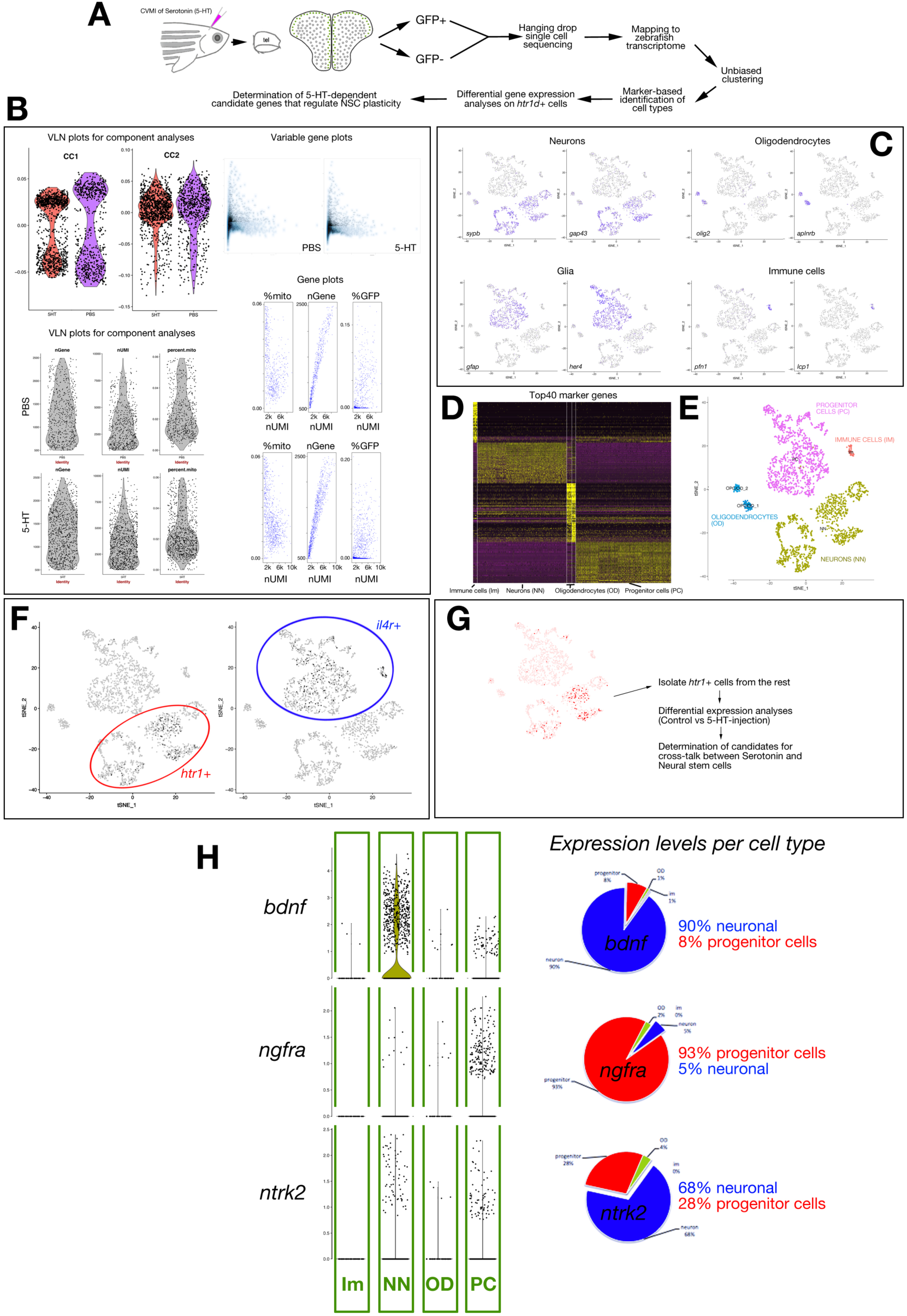
Single cell sequencing analyses of adult zebrafish telencephalon after Serotonin treatment. (A) Schematic workflow for single cell sequencing. (B) Quality control indicators of single cell sequencing data: VLN plots for principal component analyses, variable gene plots, distribution plots for number of genes (nGene), number of reads (nUMI), % of mitochondrial genes (% mito), and gene plots for % mito, nGene and %GFP (from sorted her4.1-GFP cells). (C) Primary tSNE feature plots indicating major cell clusters with canonical markers: *sypb* and *gap43* for neurons, *olig2* and *aplnrb* for oligodendrocytes, *gfap* and her4 for glia, *pfn1* and *lcp1* for immune cells. (D) Primary heat map for top 40 marker genes of neurons, glia, oligodendrocytes and immune cells. (E) Classification of major cell clusters for their identities based on markers. (F) Feature plots for *htr1* and *il4r* expression. Note that *htr1*-positive cells are neurons while *il4r*-positive cells are glia. (G) Strategy for isolating *htr1*-expressing cells from tSNE plot and subsequent differential expression analyses. (H) VLN plots for *bdnf, ngfra* and *ntrk2* in major cell types, and expression level ratios as pie charts. Related to Figure 3.

**FigureS4:**
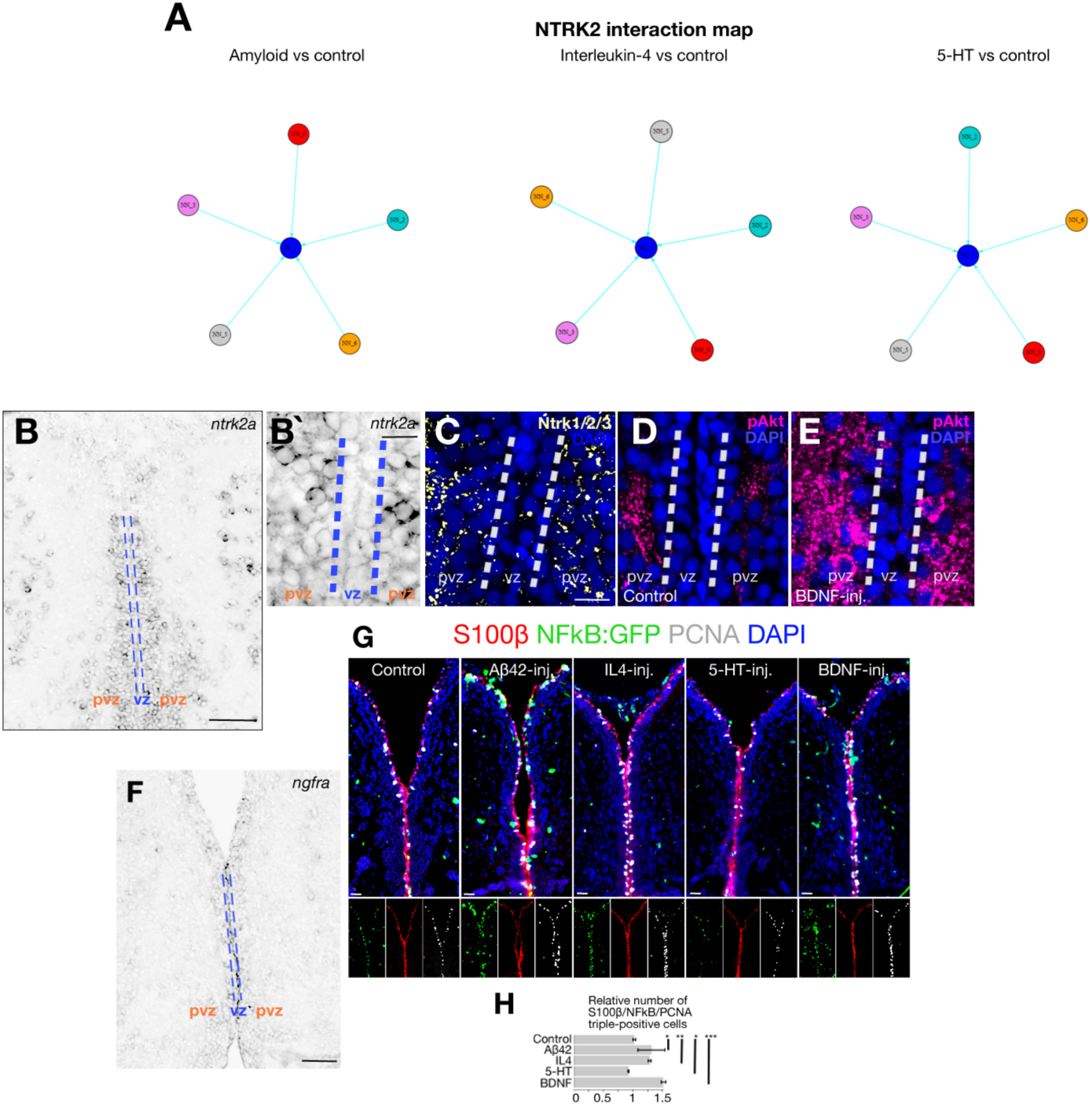
Serotonin suppresses and BDNF enhances NFkB signaling in neural stem cells in zebrafish. (A) In silico interaction map for NTRK2 in Amyloid vs control, IL4 vs control and 5-HT vs control comparisons. Black arrows: interactions unchanged with treatment, cyan arrows: interaction lost with treatment, magenta arrows: interaction gained/emerged with the treatment. (B) In situ hybridization for *ntrk2* in zebrafish brain. B’: close up image. Note the expression in periventricular zone (pvz) but not in ventricular zone (vz) that contains the neural stem cells. (C) Immunohistochemistry (IHC) for Ntrk2 protein in zebrafish brain, supporting the in situ hybridization results and presence of Ntrk2 in pvz. (D,E) IHC for pAkt in control (D) and BDNF-injected (E) brains. BDNF activates pAkt in pvz but not in vz. (F) In situ hybridization for *ngfra* in adult zebrafish telecephalon. (G) Immunohistochemistry for S100b, NfkB-driven GFP and PCNA in control, Amyloid-injected, IL4-injected, 5-HT-injected and BDNF-injected brains. Smaller panels under larger images show individual fluorescent channels. (H) Quantification of the relative number of proliferating neural stem cells that have active NFkB signaling. Scale bars equal 100 µM. Data are represented as mean ± SEM. Related to Figure 4.

**FigureS5:**
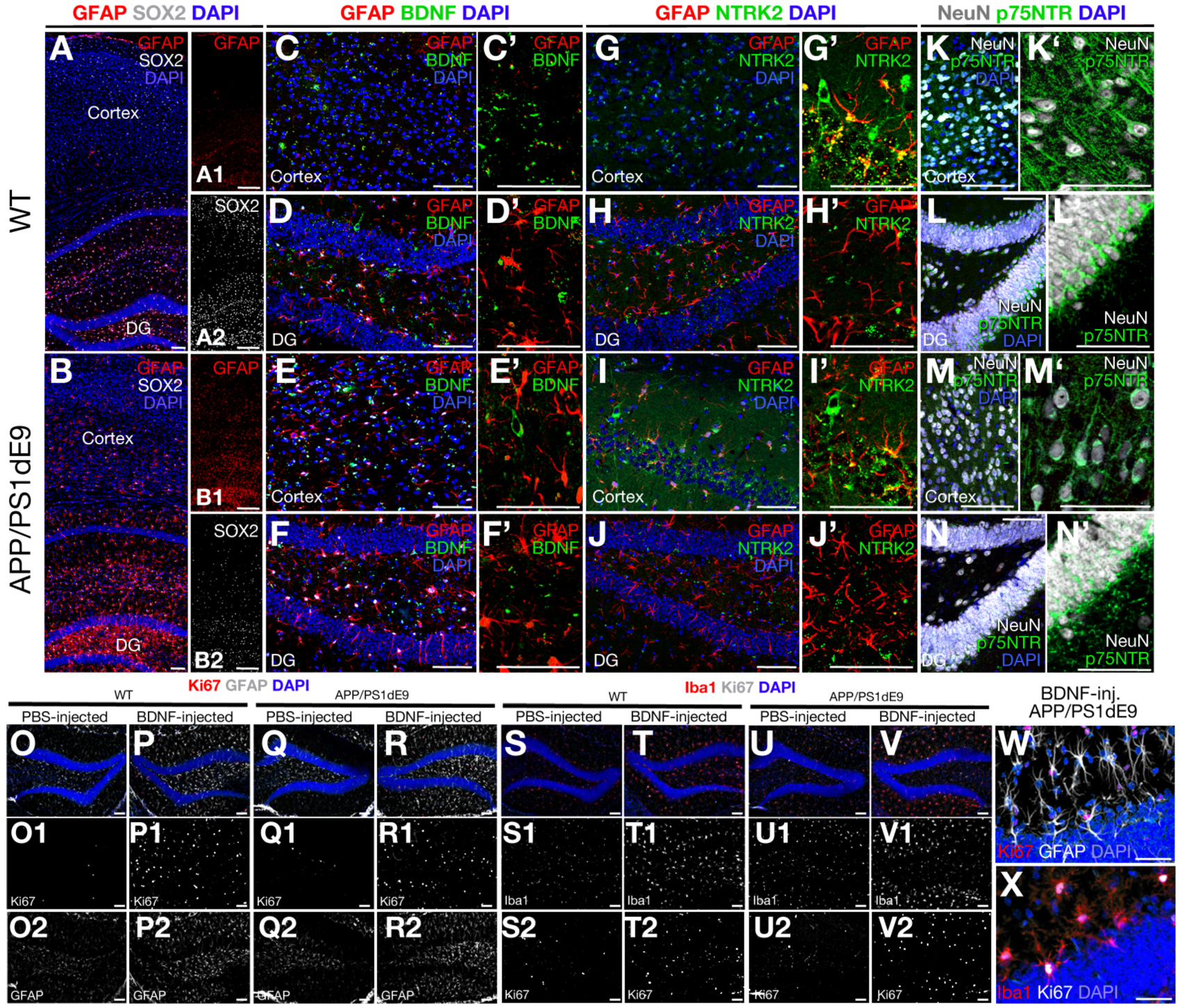
BDNF does not induce neural stem cell plasticity in mouse model of AD. (A,B) Immunohistochemistry (IHC) for GFAP and SOX2 in wild type (A) and APP/PS1dE9 mouse brains (B) at 12 months of age. (A1, A2) Single fluorescent channels of A. (B1, B2) Single fluorescent channels of B. (C-D’) IHC for GFAP and BDNF in wt mouse: C: cortex, D: dentate gyrus (DG). Primed images are higher magnification without DAPI. (E-F’) IHC for GFAP and BDNF in APP/PS1dE9 mouse: E: cortex, F: DG. Primed images are higher magnification without DAPI. (G-H’) IHC for GFAP and NTRK2 in wt mouse: G: cortex, H: DG. Primed images are higher magnification without DAPI. (I-J’) IHC for GFAP and NTRK2 in APP/PS1dE9 mouse: I: cortex, J: DG. Primed images are higher magnification without DAPI. (K-L’) IHC for NeuN and p75/NTR in wt mouse: K: cortex, L: DG. Primed images are higher magnification without DAPI. (M-N’) IHC for NeuN and p75/NTR in APP/PS1dE9 mouse: M: cortex, N: DG. Primed images are higher magnification without DAPI. (O-R2) IHC for Ki67 and GFAP in wt mouse PBS-injected hemisphere (O), wt mouse BDNF-injected hemisphere (P), APP/PS1dE9 mouse PBS-injected hemisphere (Q), APP/PS1dE9 mouse BDNF-injected hemisphere (R). (S-V2) IHC for Ki67 and Iba1 in wt mouse PBS-injected hemisphere (S), wt mouse BDNF-injected hemisphere (T), APP/PS1dE9 mouse PBS-injected hemisphere (U), APP/PS1dE9 mouse BDNF-injected hemisphere (V). (W) High magnification image of Ki67 and GFAP staining on DG of BDNF-injected hemisphere of APP/PS1dE9 mouse. GFAP cells do not proliferate after BDNF injection. (X) High magnification image of Ki67 and Iba staining on DG of BDNF-injected hemisphere of APP/PS1dE9 mouse. Most Ki67-positive cells overlap with Iba1 staining after BDNF injection. n =2 animals used for analyses. Scale bars equal 100 µM. Related to Figure 4.

**FigureS6:**
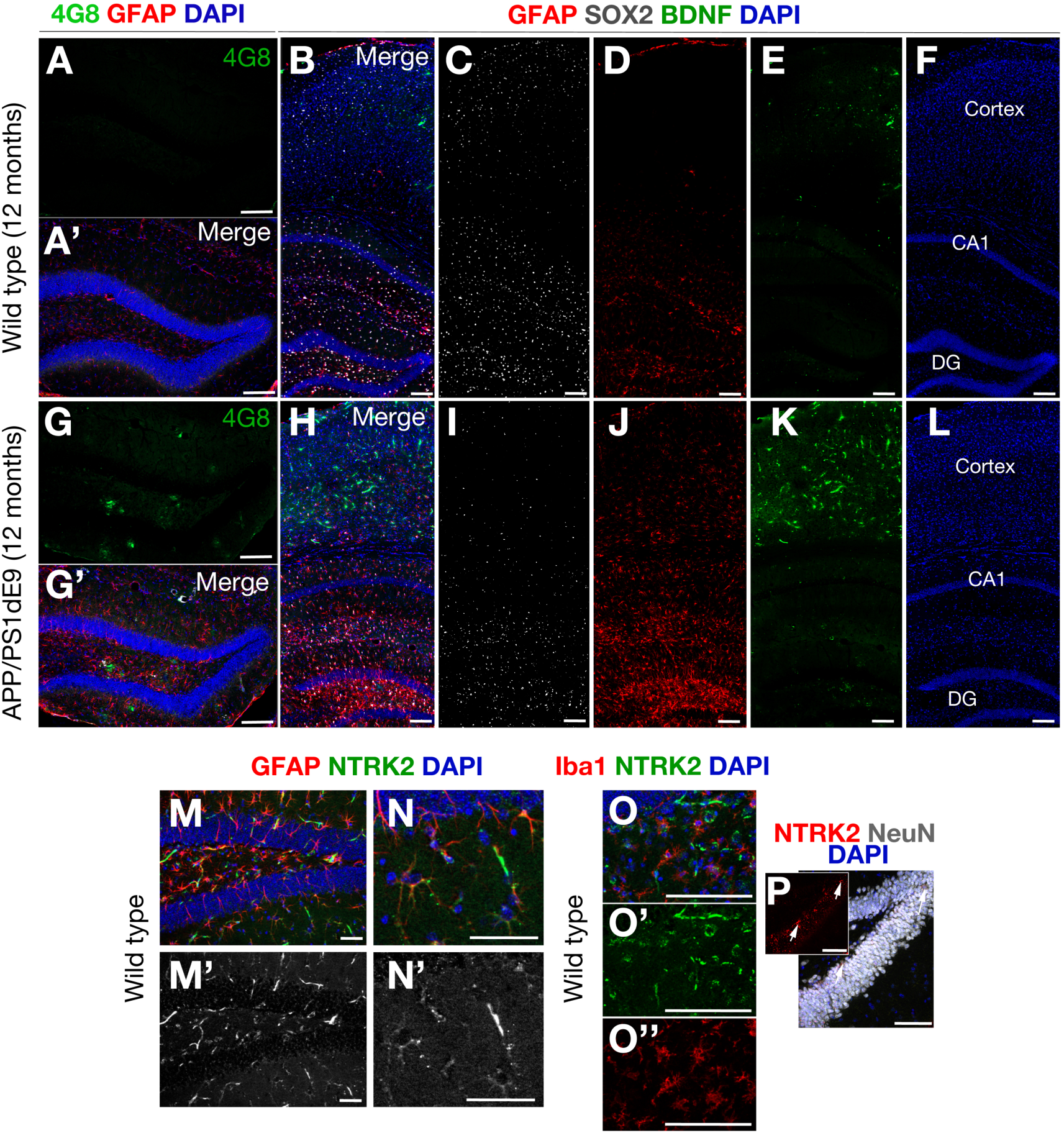
BDNF and NTRK2 expression in mouse brain. (A, A’) IHC for Amyloid plaques (4G8) in wild type mouse hippocampus. (B-F) IHC for GFAP, SOX2 and BDNF in wild type mouse brains. (G, G’) IHC for Amyloid plaques (4G8) in APP/PS1dE9 mouse hippocampus. (H-L) IHC for GFAP, SOX2 and BDNF in APP/PS1dE9 mouse brains. Mice were at the age of 12 months. (M) IHC for GFAP and NTRK2 in wild type mouse dentate gyrus. (M’) Single fluorescent channel for NTRK2. (N) High magnification of merged image. (N’) Single fluorescent channel for NTRK2 in N. (O) IHC for Iba1 and NTRK2. O’: single fluorescent channel for NTRK2 in O. O’’: single fluorescent channel for Iba1 in O. (P) IHC for NTRK2 and NeuN. Single channel in red is NTRK2. Scale bars equal 100 µM. Related to Figure 4.

**FigureS7:**
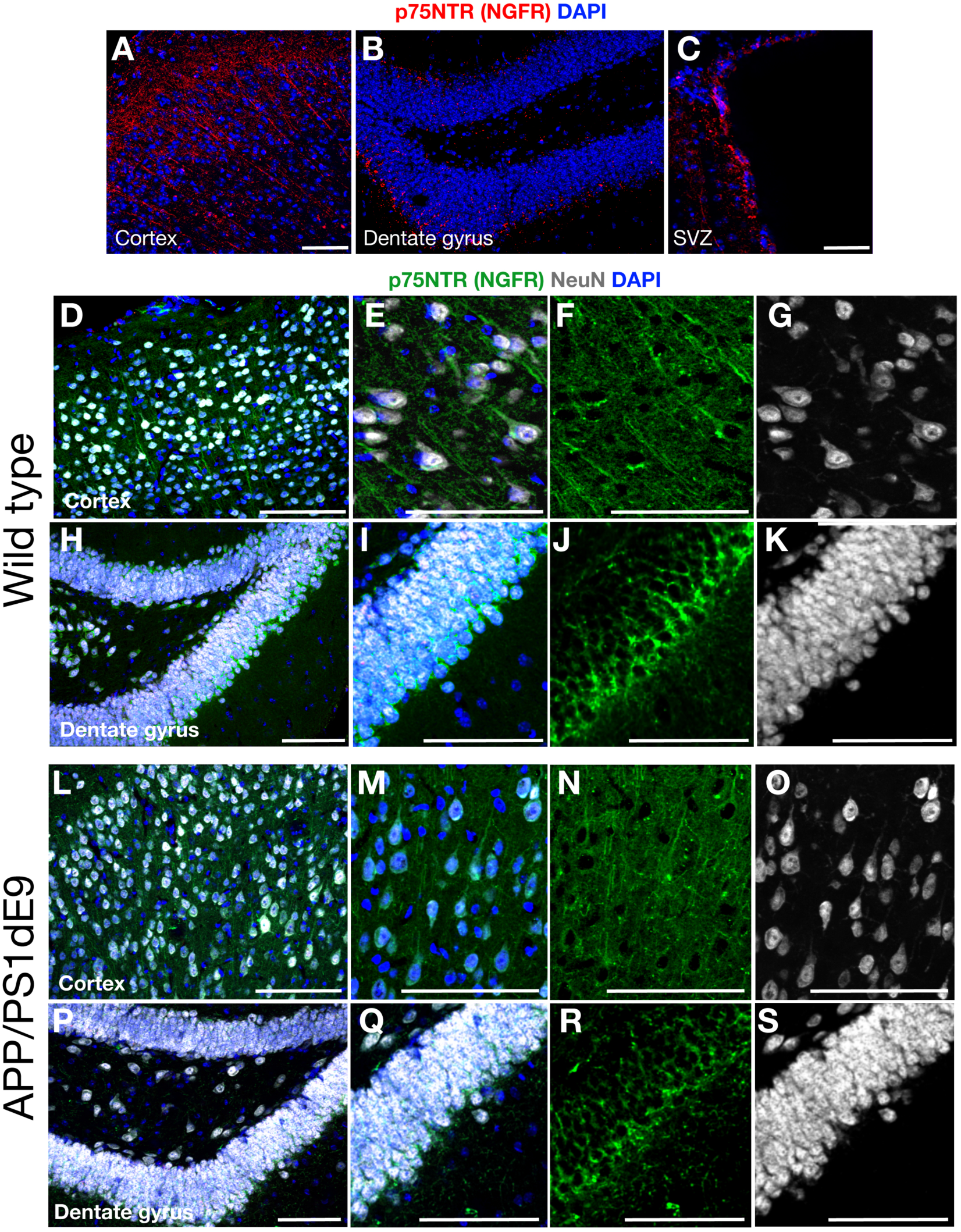
p75/NTR expression in mouse brain. (A-C) IHC for p75/NTR (NGFR) in wild type cortex (A), dentate gyrus (B) and subventricular zone, SVZ (C). (D-K) IHC for p75NTR and NeuN in cortex (D) and dentate gyrus (H). (E-G) high magnification image from D. (I-K) high magnification image from H. (L-S) IHC for p75NTR and NeuN in cortex (L) and dentate gyrus (P) of APP/PS1dE9 mouse. (M-O) High magnification image from L. (Q-S) high magnification image from P. Scale bars equal 100 µM. Related to Figure 4.

**FigureS8:**
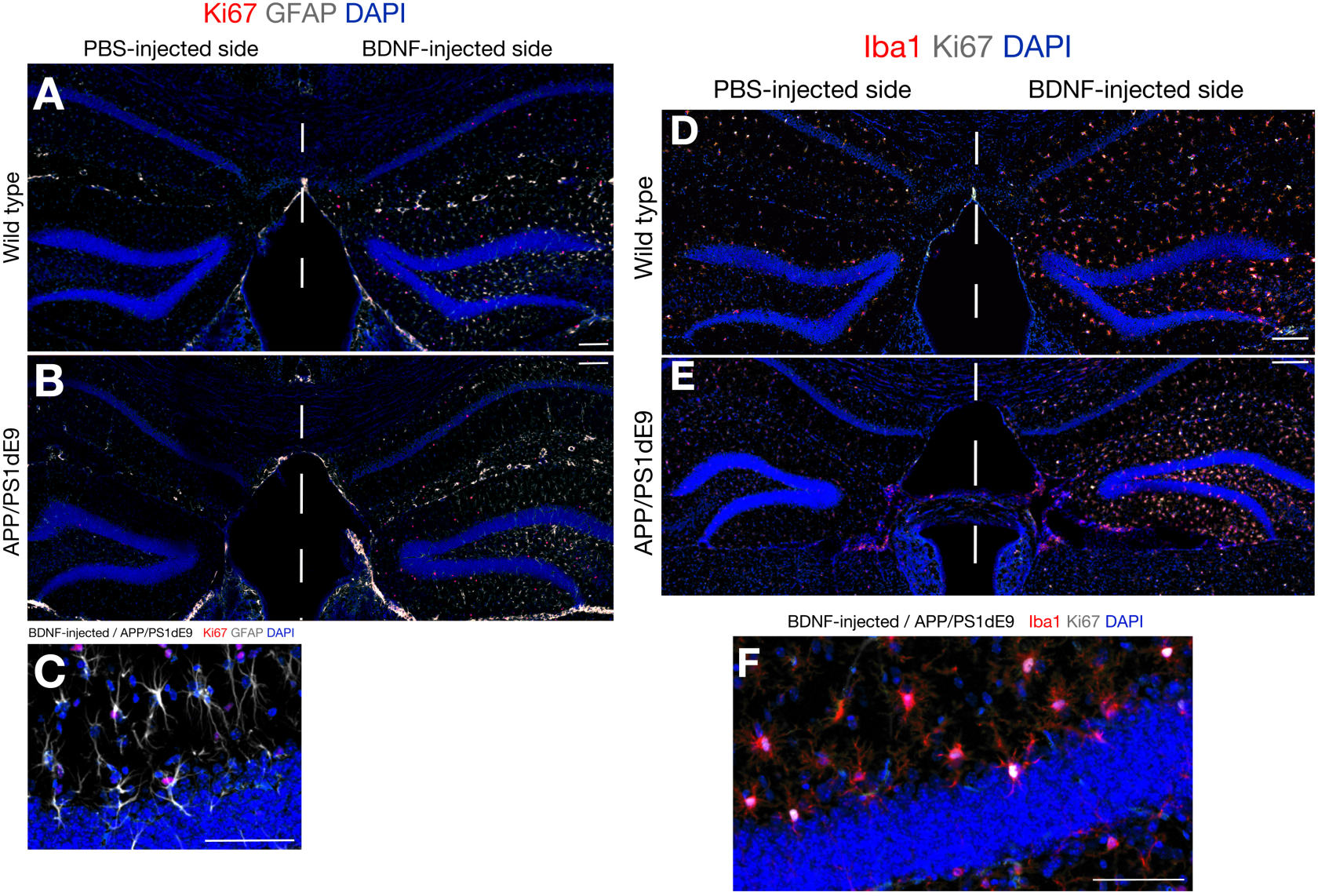
BDNF increases the proliferation of microglia but does not affect the astrocytes in the mouse brain. (A,B) IHC for Ki67 and GFAP in wild type (A) and APP/PS1dE9 (B) mouse brains at 12 months of age after injection of PBS (left hemisphere) and BDNF (right hemisphere). (C) Close up image of the dentate gyrus of a BDNF-injected side of the APP/PS1dE9 mouse brain. (D,E) IHC for Ki67 and Iba1 in wild type (D) and APP/PS1dE9 (E) mouse brains at 12 months of age after injection of PBS (left hemisphere) and BDNF (right hemisphere). (F) Close up image of the dentate gyrus of a BDNF-injected side of the APP/PS1dE9 mouse brain. Scale bars equal 100 µM. Related to Figure 4.

Dataset S1: List of differentially expressed genes determined by deep sequencing after injection of Interleukin-4 to the adult zebrafish brain.

Dataset S2: GO term and KEGG pathway analyses for transcriptomics changes in adult zebrafish brain after Interleukin-4

Dataset S3: Single cell sequencing quality control datasets: histograms, VLN plots, tSNE plots, heat maps

Dataset S4: Differentially expressed genes in *htr1*+ cells after single cell sequencing

Dataset S5: Differentially expressed genes in *il4r.1*+ cells after single cell sequencing

Dataset S6: Differentially expressed genes in *ngfra*+ cells after single cell sequencing

